# DNA catenation is essential for Sister Chromatid Cohesion

**DOI:** 10.64898/2026.07.16.738924

**Authors:** Aditi Kaushik, Alba A Fraile, Vanessa Viera, Alejandro M Martin, Andres H Maduro, James Collier, Toyonori Sakata, Jumpei Fukute, Katsuhiko Shirahige, Kristian Jeppsson, Raquel A Oliveira, Joaquim Roca, Madhusudhan Srinivasan

## Abstract

From S phase until anaphase, sister chromatids remain physically linked to ensure accurate chromosome segregation. This linkage, called sister chromatid cohesion, must withstand the pulling forces generated by the mitotic spindle and is thought to require entrapment of sister DNAs inside the ring-shaped cohesin complex. DNA catenations that arise naturally during replication could also tether sister chromatids, but whether they contribute to cohesion has remained unresolved. Here, using budding yeast, we show that cohesin-mediated sister DNA entrapment alone cannot withstand spindle-generated pulling forces and that robust sister chromatid cohesion requires DNA catenation. Selective removal of DNA catenations causes catastrophic cohesion loss, delays chromosome biorientation and increases chromosome mis-segregation, even when cohesin rings remain intact. Importantly, DNA catenation similarly underpins force-resistant cohesion in metazoan chromosomes. Our findings fundamentally redefine the physical basis of sister chromatid cohesion by establishing DNA catenation as an evolutionarily conserved component of the force-resistant linkage between sister chromatids that enables accurate chromosome segregation.

## Introduction

The continuity of eukaryotic life relies on accurate genome segregation during mitosis and meiosis, often long after DNA replication. During both processes, sister chromatids must remain physically linked from when they are replicated in S phase until their kinetochores achieve biorientation in mitosis after attachment to microtubules from opposite spindle poles. This linkage, known as sister chromatid cohesion, prevents premature separation and generates the tension needed for stable spindle attachment ^1^. Once all chromosomes are bioriented, the anaphase-promoting complex (APC/C) is activated ^2^, triggering destruction of cohesion and the ensuing disjunction of sisters to opposite poles ^3^.

The earliest model for cohesion involved the intertwining of sister DNAs or catenations of sister DNAs arising as a natural by-product of DNA replication ^4^. Though conceptually elegant, it could not explain how catenations resist being displaced toward chromosome ends by spindle forces or how they avoid premature resolution by Topoisomerase II (Topo II). Indeed, the discovery that small circular mini-chromosomes in yeast are largely de-catenated even before the cells embark on mitosis led to the search for an alternative mechanism ^5^, which resulted in the identification of a ring-shaped protein complex cohesin ^6,7^.

Cohesin belongs to the highly conserved structural maintenance of chromosomes (SMC)-kleisin family of ATPases ^8^, a group of proteins capable of capturing and processively extruding DNA loops to organize the genomes of most cells ^9–11^. In eukaryotes, cohesin structures the interphase genome^10^ while its paralog condensin drives mitotic chromosome assembly ^12^. What distinguishes cohesin from other SMC complexes is its ability to topologically entrap sister DNAs within its ring structure ^13^, a property fundamental for cohesion ^14^. This entrapment of sister DNAs persists from S phase until anaphase, when APC/C-activated separase cleaves cohesin rings, thus triggering sister chromatid disjunction ^3^.

The demonstration that cohesin can topologically entrap sister DNAs independently of DNA catenation^13^, has fostered the prevailing view that cohesin-mediated entrapment is both necessary and sufficient for sister chromatid cohesion. This ring model^15^ provides the most comprehensive explanation of cohesion to date and is supported by a vast body of experimental evidence. However, it also carries an implicit assumption that has never been directly tested: that cohesin rings by themselves are capable of resisting the pulling forces generated by the mitotic spindle.

Curiously, cohesin has also been observed to stabilise residual catenations between sister chromatids ^16–18^. Whether these catenations are merely vestigial remnants of replication that cohesin passively preserves, or whether they play an essential functional role in maintaining cohesion under spindle tension, has never been tested. Here, by measuring chromosomal catenation at centromeric and peri-centric regions and selectively eliminating catenations, we show that sister chromatid cohesion critically depends on persistent DNA catenation.

## Results

### Catenations preserved by cohesin accumulate at peri-centromeres in mitosis

To determine whether DNA catenations contribute to sister chromatid cohesion, we first measured their abundance within centromeric and pericentromeric regions of budding yeast chromosomes, where cohesin is enriched^19^. We engineered loxP sites flanking either the centromere of Chromosome III (CEN III) or a neighbouring pericentric region (Fig. 1A) ^19^. Induction of Cre recombinase in mitotically arrested cells treated with nocodazole excised CEN III in approximately 75% of cells (Fig. S1A,B). Genomic DNA was isolated and the excised DNA circles analysed by Southern blotting using CEN- or peri-CEN-specific probes (Fig. 1A). Because nocodazole depolymerizes spindle microtubules, these measurements were made in metaphase-arrested cells in the absence of spindle forces. Under these conditions, the excised DNA circles migrated as multiple topological forms (Fig. 1A). Nicking converted these into two species: monomeric circles and catenated sister DNA circles, which were completely resolved into monomers by *E. coli* Topo IV (Fig. 1A, Fig. S1C). Using this assay, we found that approximately 30% of excised centromeric circles from wild-type mitotic cells were catenated, corresponding to roughly one catenation per 10 kb of centromeric DNA (Fig. 1B). Depletion of the cohesin subunit Scc1 markedly reduced these catenations (Fig. 1B, Fig. S1D, E), confirming that cohesin preserves sister DNA catenations^16,17^.

**Fig. 1.**
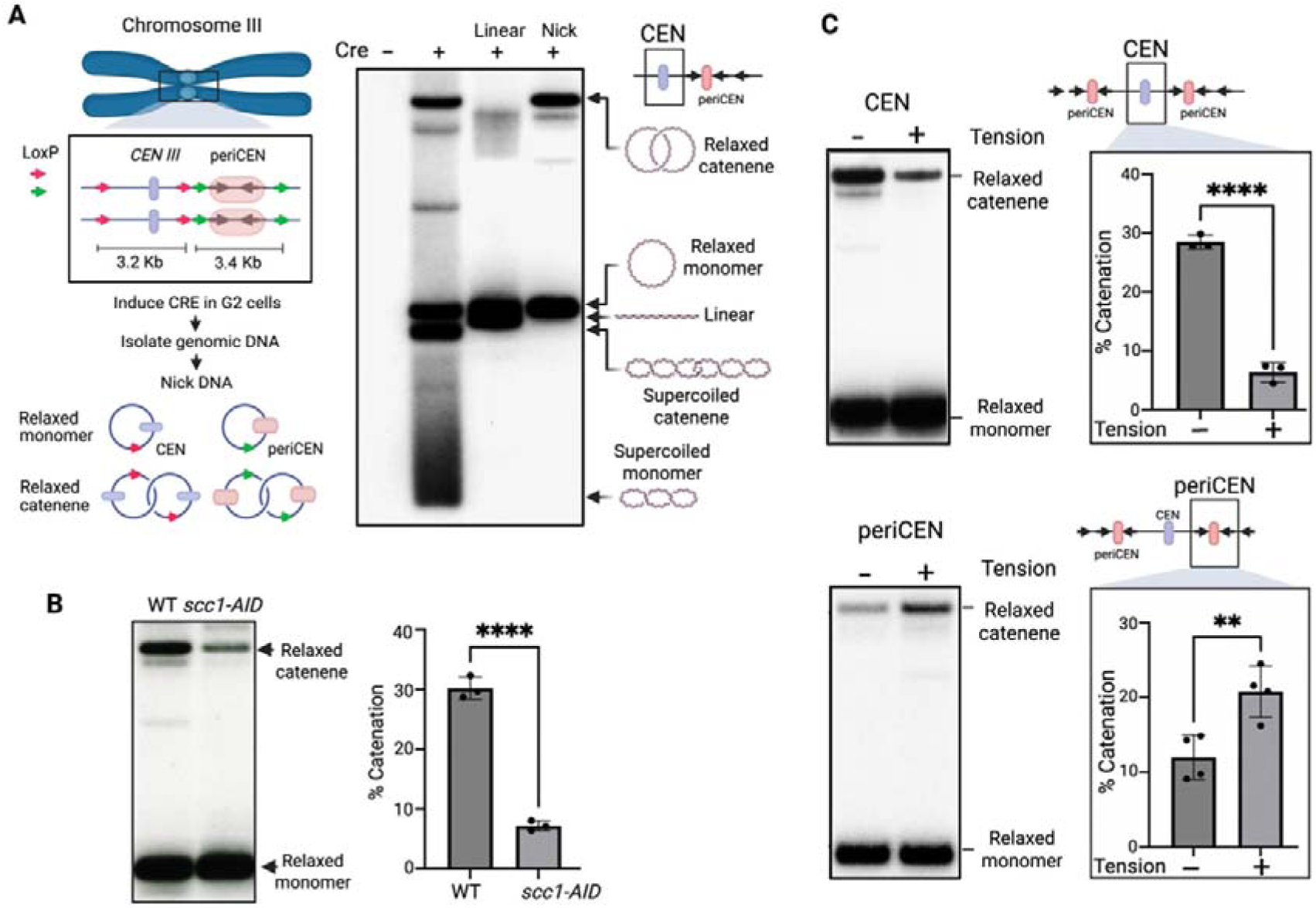
Cohesin preserves DNA catenations that accumulate at peri-centromeres under spindle tension. (A) Assay for measuring centromeric (CEN) and pericentromeric (peri-CEN) DNA catenation. LoxP sites were introduced flanking either the centromere III (CEN III) or a neighbouring pericentromeric region of chromosome III. Typically, cells were arrested in mitosis and Cre recombinase was induced as described in Fig. S1A and *Methods*. Genomic DNA was isolated from fixed cells, linearized with HindIII or nicked with Nb.BsmI, and analysed by agarose gel electrophoresis and Southern blotting using CEN or peri-CEN probes. (B) Cohesin preserves centromeric DNA catenations. Wild-type (29532) and SCC1-AID (30245) cells were synchronised in G1, depleted of Scc1 by auxin addition, and released into mitotic arrest in the presence of nocodazole and auxin (Fig. S1D,E; Methods). CEN III was excised, genomic DNA was isolated, nicked and analysed by Southern blotting. Right, quantification of DNA catenation from three independent biological experiments. Data are mean ± s.d. (C) Spindle tension redistributes DNA catenations from centromeres to pericentromeres. DNA catenation at CEN (30411) and peri-CEN (30904) regions was measured in metaphase-arrested cells in the absence (nocodazole) or presence (Cdc20 depletion) of spindle tension (Fig. S1G; Methods). Representative Southern blots are shown together with quantification from three independent biological experiments. Data are mean ± s.d.

Having established that cohesin preserves centromeric DNA catenation in the absence of spindle forces, we next asked how spindle tension influences their distribution. Because spindle forces redistribute cohesin from centromeres (CEN) to pericentromeric (peri-CEN) regions during metaphase^19^ (Fig. S1F), cells were arrested in metaphase by depletion of the APC co-activator Cdc20, either in the presence or absence of nocodazole, and CEN and peri-CEN regions were excised and analysed. Strikingly, spindle tension caused centromeric catenations to decrease, accompanied by a corresponding increase in pericentromeric catenations (Fig. 1C, Fig. S1G), indicating that DNA catenations redistribute from CEN to peri-CEN under tension. Remarkably, despite persistent spindle-generated pulling forces, DNA catenations remained confined to pericentromeric regions rather than being displaced further along chromosome arms. This retention suggests that specific features of the peri-centromere act as barriers that spatially confine both cohesin and DNA catenations under spindle tension.

### DNA catenation alone cannot generate a force-resistant cohesion

The persistence of DNA catenations within pericentromeric regions raised the possibility that they make a functional contribution to cohesion. Whether DNA catenation alone can generate cohesion capable of resisting spindle-generated pulling forces has never been directly tested. Addressing this question is challenging because cohesin is required both to establish cohesion and to preserve DNA catenations. Simply removing cohesin therefore eliminates both mechanisms simultaneously, making it impossible to determine the contribution of DNA catenation alone.

To overcome this problem, we first assessed sister chromatid cohesion in mitotically arrested cells lacking spindle forces by monitoring GFP-marked peri-centromere on chromosome IV (Fig. 1D, Fig. S1H). Inactivation of cohesin using the temperature-sensitive *scc1-73* allele led to separation of sister loci in approximately 75% of cells (Fig. 2A, Fig. S1H), consistent with previous reports ^6,7^. This loss of cohesion could result either from loss of sister DNA entrapment inside cohesion rings or from the accompanying loss of DNA catenations normally preserved by cohesin. We therefore asked whether preventing Topo II-mediated decatenation could restore cohesion. Simultaneous inactivation of Topo II using the temperature-sensitive *top2-4* ^20^ allele markedly suppressed sister chromatid separation (Fig. 2A), providing independent functional evidence that DNA catenations persist on mitotic chromosomes. These catenations can partially compensate for cohesin loss by maintaining sister chromatid proximity in the absence of spindle forces (Fig. 2A).

**Fig. 2.**
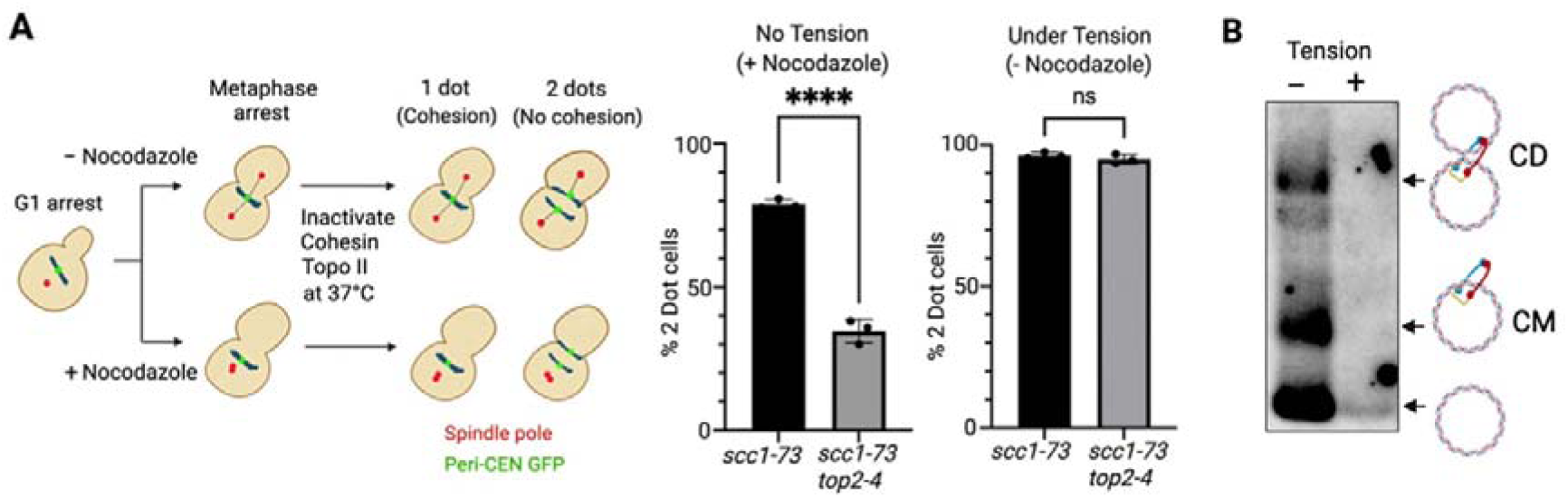
Neither cohesin-mediated DNA entrapment nor DNA catenation alone is sufficient for cohesion. (A) DNA catenation alone cannot maintain cohesion under spindle tension. Cohesion at a GFP-marked pericentromeric locus on chromosome IV was assayed in *scc1-73* (31001) and *scc1-73 top2-4* (31044) cells. Cells were synchronised in G1, released into metaphase in the presence or absence of nocodazole, and cohesin and/or Topo II were inactivated by shifting to 37°C (Fig. S1H; Methods). Cohesion was scored cytologically as one GFP focus (cohesed sister chromatids) or two GFP foci (separated sister chromatids). At least 100 metaphase cells were analysed per experiment. Graphs show mean ± s.d. from three independent biological experiments. **(B) Cohesin-mediated sister DNA entrapment is not maintained under spindle tension in the absence of DNA catenation.** Topological co-entrapment of sister DNAs by cohesin was measured using the 6C assay. Cells carrying the 2.3 kb mini-chromosome (32191) were synchronised in G1 and released into metaphase by Cdc20 depletion in the presence or absence of spindle tension (Fig. S2A; Methods). All three cohesin ring interfaces were chemically crosslinked in vivo before cohesin immunoprecipitation under denaturing conditions and Southern blot analysis of the associated mini-chromosome DNA. CD, sister mini-chromosome co-entrapment within a single cohesin ring; CM, entrapment of a single mini-chromosome.

The crucial question, however, was whether DNA catenation alone could also sustain cohesion under spindle tension. To test this, we repeated the same experiment in metaphase-arrested cells with bipolar spindles established through Cdc20 depletion. Strikingly, when spindle generated pulling forces were established, sister loci separated in approximately 95% of cells despite the persistence of DNA catenations (Fig. 2A). Thus, although DNA catenations can passively tether sister chromatids in the absence of spindle forces, they cannot withstand spindle-generated pulling forces without cohesin, most likely because tension displaces them towards chromosome ends (Fig. S1I). DNA catenation alone therefore cannot provide the force-resistant linkage required to maintain sister chromatid cohesion during mitosis.

### Cohesin rings cannot withstand spindle forces without catenations

Our finding that DNA catenation alone cannot sustain cohesion under spindle tension raised the reciprocal question: is cohesin-mediated sister DNA entrapment alone sufficient to resist spindle-generated pulling forces? Previous work using a small 2.3 kb mini-chromosome, which carries negligible DNA catenation^21^, showed that cohesin rings can co-entrap sister DNAs independently of catenations^13^. Whether such cohesin-mediated entrapment can withstand spindle tension during mitosis, however, has never been tested.

To address this, we used the 6C assay^14^, which measures topological DNA entrapment by cohesin. In this assay, all three ring interfaces are chemically crosslinked in vivo, permanently sealing the cohesin ring. Cohesin is then immunoprecipitated, denatured with SDS, and the associated mini-chromosome DNA detected by Southern blotting. Because the rings are covalently closed, any enclosed DNA remains trapped inside cohesin rings, allowing identification of single or sister mini-chromosomes captured by cohesin.

Cells were arrested in metaphase either in the presence or absence of spindle forces and analysed by the 6C assay (Fig. S2A). Consistent with previous observations ^14^, cohesin robustly co-entrapped sister mini-chromosomes in the absence of spindle tension (Fig. 2B). Strikingly, however, no stable sister mini-chromosome co-entrapment was detected when spindle forces were applied (Fig. 2B), despite cohesin being immunoprecipitated and chemically circularised with equal efficiency under both conditions (Fig. S2B). Thus, in the absence of DNA catenation, cohesin-mediated sister DNA entrapment alone cannot withstand spindle-generated pulling forces. Together with our finding that DNA catenation alone cannot sustain cohesion under spindle tension, these observations demonstrate that neither cohesin-mediated sister DNA entrapment nor DNA catenation alone is sufficient to generate force-resistant cohesion. Rather, the two topological mechanisms must act together to generate the force-resistant linkage that resists spindle pulling forces.

### Catenations are necessary for cohesion to withstand spindle forces

Our finding that neither DNA catenation nor cohesin-mediated sister DNA entrapment alone can withstand spindle forces suggested that force-resistant sister chromatid cohesion emerges from their coordinated action. A key prediction of this model is that cohesin should no longer maintain cohesion if DNA catenations are selectively removed. Testing this prediction requires eliminating DNA catenations while preserving cohesin-mediated sister DNA entrapment.

To acheive this, we expressed a highly active topoisomerase II from *Chlorella* virus (PBCV-Topo II) (Fig. S2D), which efficiently resolves DNA catenations^21^. Before using PBCV-Topo II to test cohesion, we first established that it selectively removes DNA catenations without disrupting cohesin function. In nocodazole-arrested cells carrying a large circular mini-chromosome, PBCV-Topo II expression completely eliminated catenated DNA species (Fig. S2E), demonstrating that it overcomes cohesin-mediated protection of DNA catenations. Differential sedimentation of mini-chromosome dimers showed that cohesin-mediated sister DNA co-entrapment remained unaffected (Fig. S2E), while pulsed-field gel electrophoresis revealed no evidence of chromosome breakage (Fig. S2F). Together, these controls establish that PBCV-Topo II selectively removes DNA catenations while preserving cohesin-mediated sister DNA entrapment and overall chromosome integrity.

Having established a means to selectively eliminate DNA catenations, we next tested whether cohesin alone can maintain cohesion under spindle tension. Wild-type cells were arrested in metaphase by Cdc20 depletion, PBCV-Topo II was induced, and cohesion was monitored at a pericentromeric locus on chromosome IV (Fig. S2G). Strikingly, selective removal of DNA catenations triggered catastrophic sister chromatid separation despite the continued presence of cohesin (Fig. 3A). This defect was completely suppressed by depolymerizing spindle microtubules with nocodazole (Fig. 3A), demonstrating that cohesion fails specifically when chromosomes are subjected to spindle-generated pulling forces. Thus, although cohesin alone can maintain sister chromatid proximity in the absence of tension, DNA catenation is required to generate the force-resistant linkage that maintains cohesion under spindle tension.

**Fig. 3.**
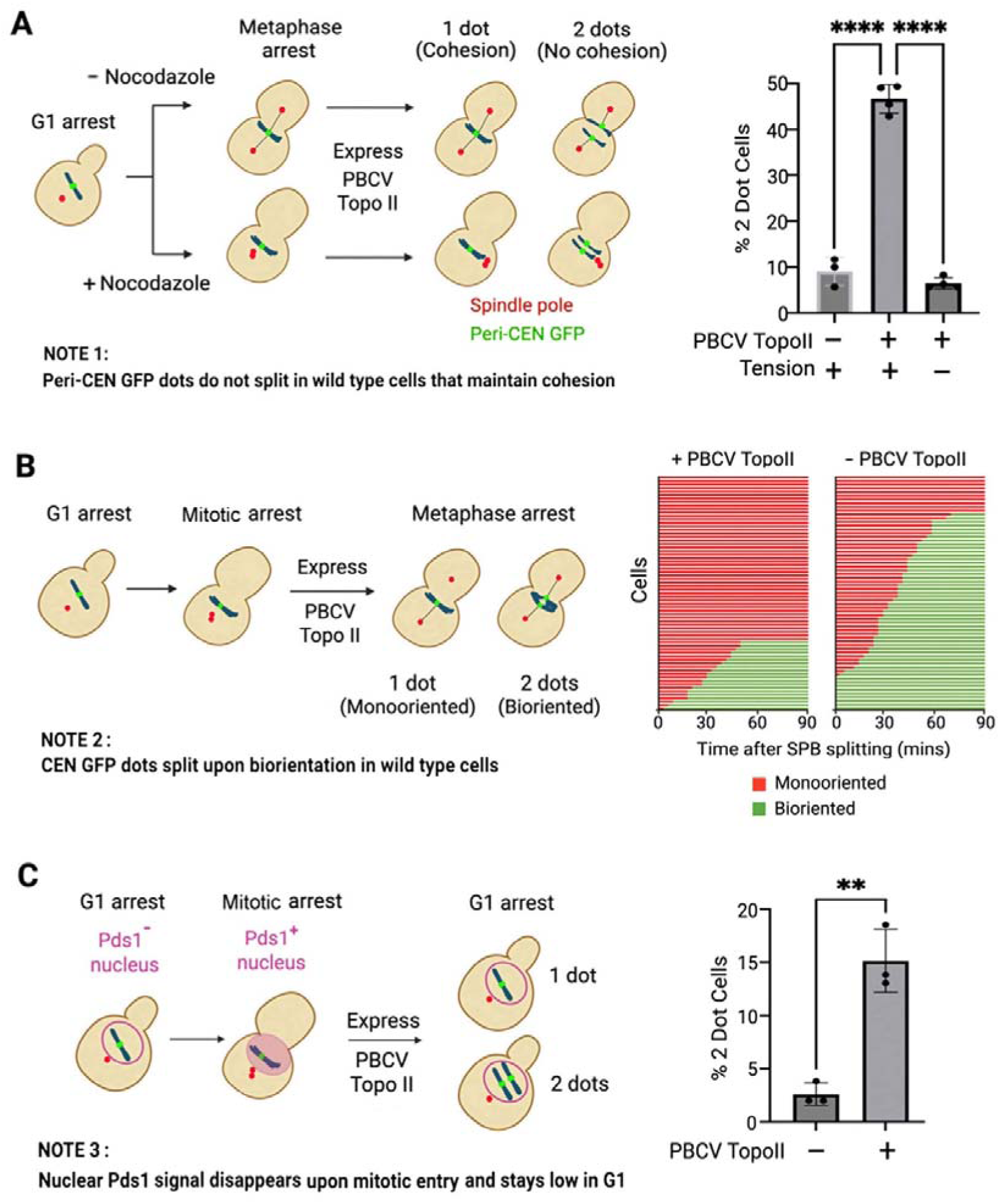
Selective removal of DNA catenations compromises force-resistant cohesion, chromosome biorientation and chromosome segregation. (A) Selective removal of DNA catenations abolishes force-resistant cohesion. Cohesion at a GFP-marked pericentromeric locus on chromosome IV was assayed in cells carrying either an empty vector (30484) or GAL1-PBCV-TOPII (30548). Cells were synchronised in G1, released into metaphase in the presence or absence of nocodazole, and PBCV-Topo II expression induced following metaphase arrest (Fig. S2G; Methods). Cohesion was scored cytologically as one GFP focus (cohesed sister chromatids) or two GFP foci (separated sister chromatids). At least 100 metaphase cells were analysed per experiment. Graphs show mean ± s.d. from three independent biological experiments. **(B) Selective removal of DNA catenations delays chromosome biorientation.** Cells carrying either an empty vector (32073) or GAL1-PBCV-TOPII (32074) were synchronised in G1, arrested in mitosis with benomyl and nocodazole, induced for PBCV-Topo II expression, and released into metaphase (Fig. S2H; Methods). Chromosome biorientation was monitored by live-cell imaging of GFP-marked CEN IV. Biorientation was scored by separation of sister CEN IV loci following spindle pole body separation. Individual cells (n = 60) were followed for 90 min. **(C) Selective removal of DNA catenations compromises chromosome segregation.** Cells carrying either an empty vector (32020) or GAL1-PBCV-TOPII (32021) were synchronised in mitosis with benomyl and nocodazole, induced for PBCV-Topo II expression, and released into the subsequent cell cycle. Cells entering G1 were identified by loss of nuclear Pds1-Myc18 staining, and segregation of chromosome IV was assessed by monitoring GFP-marked CEN IV. At least 100 Pds1-negative G1 cells were analysed per experiment. Graphs show mean ± s.d. from three independent biological experiments.

### Loss of catenations compromises chromosome segregation fidelity

If DNA catenation contributes to the force-resistant linkage that maintains sister chromatid cohesion, its selective removal should compromise chromosome segregation despite intact cohesin rings. To test this prediction, we first examined chromosome biorientation, the process by which sister kinetochores establish attachment to spindle microtubules emanating from opposite spindle poles. Cells were arrested in mitosis without spindle forces using nocodazole, DNA catenations were selectively removed by induction of PBCV-Topo II, and cells were then released into metaphase to allow spindle formation. Biorientation of chromosome IV was monitored in live cells using a fluorescently marked centromere IV. Under these conditions, where cohesin-mediated sister DNA entrapment remains intact but DNA catenations have been removed, chromosome IV exhibited a pronounced delay in achieving bipolar attachment (Fig. 3B, Fig. S2H), with approximately 75% of cells failing to biorient during the course of the experiment (Fig. 3B). Thus, DNA catenation is required for efficient chromosome biorientation.

We next asked whether loss of DNA catenation compromises chromosome segregation itself. DNA catenations were selectively removed from mitotically arrested cells by induction of PBCV-Topo II in the presence of nocodazole before cells were released to complete mitosis and arrested in the subsequent G1 phase. Selective removal of DNA catenations resulted in a marked increase in chromosome mis-segregation, manifested by unequal inheritance of fluorescently marked chromosome IV (Fig. 3C). Thus, selective removal of DNA catenations compromises both chromosome biorientation and chromosome segregation despite intact cohesin rings.

These findings demonstrate that the force-resistant linkage provided by DNA catenation is essential not only for maintaining sister chromatid cohesion under spindle tension but also for ensuring faithful chromosome segregation.

### DNA catenation underpins force-resistant cohesion in metazoan chromosomes

Having established that DNA catenation is required for force-resistant sister chromatid cohesion in budding yeast, we next asked whether this represents a conserved feature of eukaryotic chromosomes. To address this, we took advantage of a Drosophila embryo system that provides a particularly stringent test of cohesion. Previous work showed that metaphase chromosomes in syncytial embryos contain an apparent excess of cohesin, such that acute removal of approximately half of the chromosome-bound cohesin has no detectable effect on sister chromatid cohesion despite persistent spindle pulling forces^22^. We reasoned that if DNA catenation similarly contributes to the force-resistant linkage in metazoan chromosomes, its role should become apparent under precisely these conditions, where cohesin has been substantially reduced but chromosomes remain fully cohesive.

Consistent with previous observations, acute removal of approximately 50% of cohesin by TEV protease injection caused no detectable cohesion fatigue, a phenomenon in which persistent spindle pulling forces gradually overcome sister chromatid cohesion during metaphase arrest, over the 15–25 minute observation period (Fig. 4). In striking contrast, simultaneous injection of TEV protease together with PBCV-Topo II to promote DNA decatenation resulted in a significant increase in embryos displaying cohesion fatigue, manifested by precocious sister chromatid separation (Fig. 4). These observations are complemented by an accompanying study by Cui. R et al, demonstrating that DNA catenation similarly underpins force-resistant sister chromatid cohesion in human cells.

**Fig. 4.**
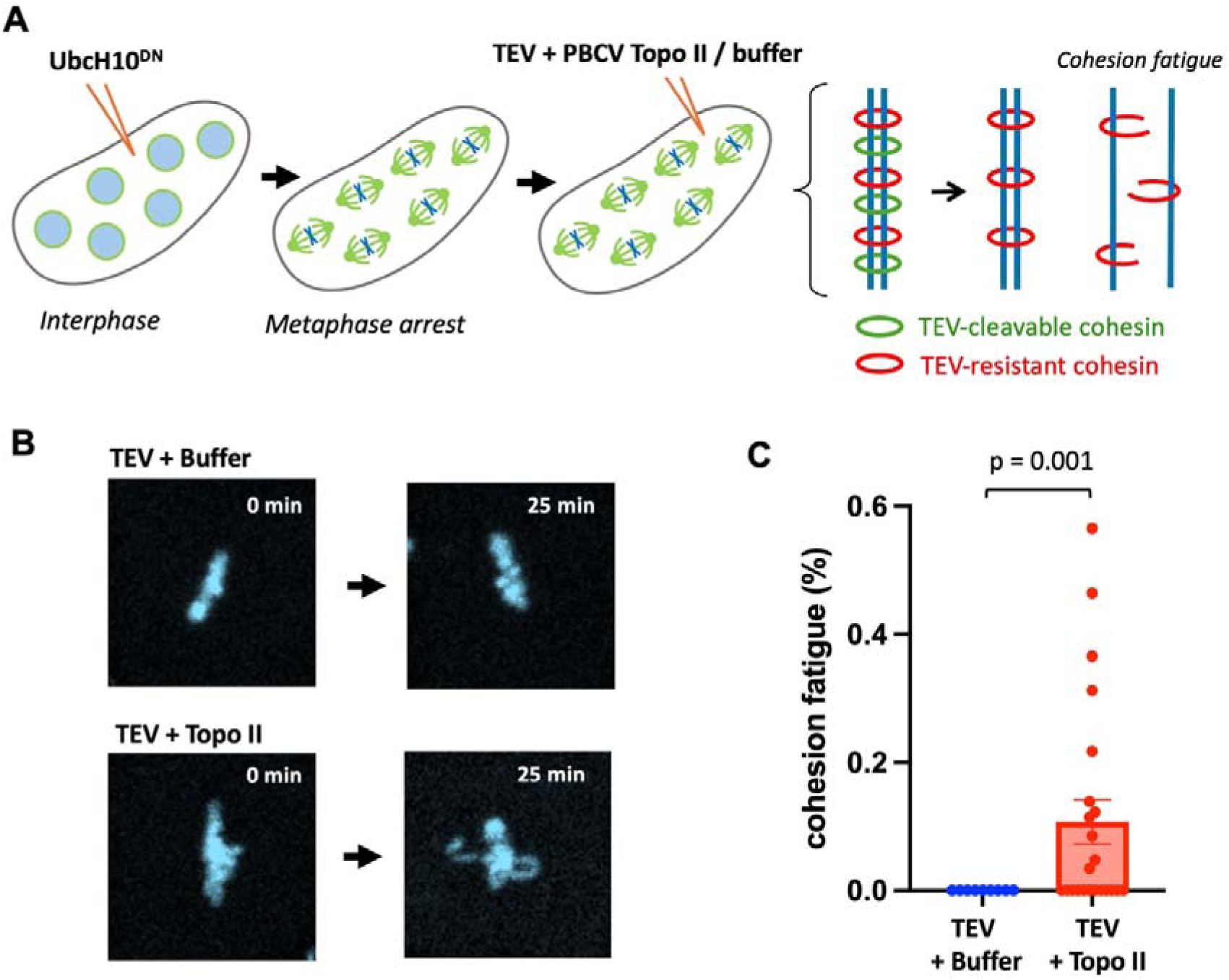
DNA catenation underpins force-resistant cohesion in Drosophila chromosomes. **(A)** Experimental scheme. Syncytial embryos expressing **UbcH10DN** were injected with TEV protease to remove the TEV-cleavable pool of cohesin (∼50% of chromosome-bound cohesin) together with either buffer or PBCV-Topo II. Embryos were maintained in metaphase arrest and chromosome behaviour monitored by live imaging. Green rings denote TEV-cleavable cohesin and red rings TEV-resistant cohesin. **(B)** Representative images of metaphase chromosomes immediately before (0 min) and 25 min after injection with TEV protease plus buffer (top) or TEV protease plus PBCV-Topo II (bottom). Images illustrate cohesion fatigue following selective removal of DNA catenations. **(C)** Quantification of cohesion fatigue. Each point represents the percentage of metaphases displaying cohesion fatigue within an individual embryo scored 15–25 min after the second injection (TEV + buffer, *n* = 8 embryos; TEV + PBCV-Topo II, *n* = 23 embryos). Bars represent mean ± s.e.m. Statistical significance was determined using the Wilcoxon rank-sum test.

Thus, DNA catenation is an evolutionarily conserved component of the force-resistant linkage that maintains sister chromatid cohesion.

### Cohesin protects catenations independently of sister DNA entrapment

Having established that DNA catenation is essential for sister chromatid cohesion and that cohesin preserves these catenations, we next sought to understand how cohesin protects them from Topo II-mediated resolution. We considered the simple possibility that cohesin protects DNA catenations simply by co-entrapping sister DNAs. Sister DNA co-entrapment might either shield DNA crossover points from Topo II-mediated resolution or maintain sister DNAs in close proximity, thereby promoting continual re-catenation and sustaining a dynamic equilibrium of DNA crossings (Fig. S3A). These mechanisms are not mutually exclusive and could operate together.

To determine whether sister DNA co-entrapment is required for catenation preservation, we analysed the consequences of depleting Pds5, a cohesin-associated factor required for stable sister DNA co-entrapment but dispensable for entrapment of individual DNA molecules^14^ and for loop extrusion ^23^. *PDS5-AID* cells were arrested in G1 using α-factor and then released into mitosis in the presence of nocodazole and auxin to induce Pds5 degradation. As expected, loss of Pds5 abolished sister DNA co-entrapment (Fig. S3D)^14^. Strikingly, however, Pds5 depletion had little or no effect on centromeric DNA catenation (Fig. 5A, Fig. S3B) and, if anything, increased pericentromeric DNA catenation (Fig. 5B). Thus, sister DNA co-entrapment is not required to preserve DNA catenations, at least in the absence of spindle forces. Instead, DNA catenations are protected by a distinct, non-cohesive pool of cohesin.

**Fig. 5.**
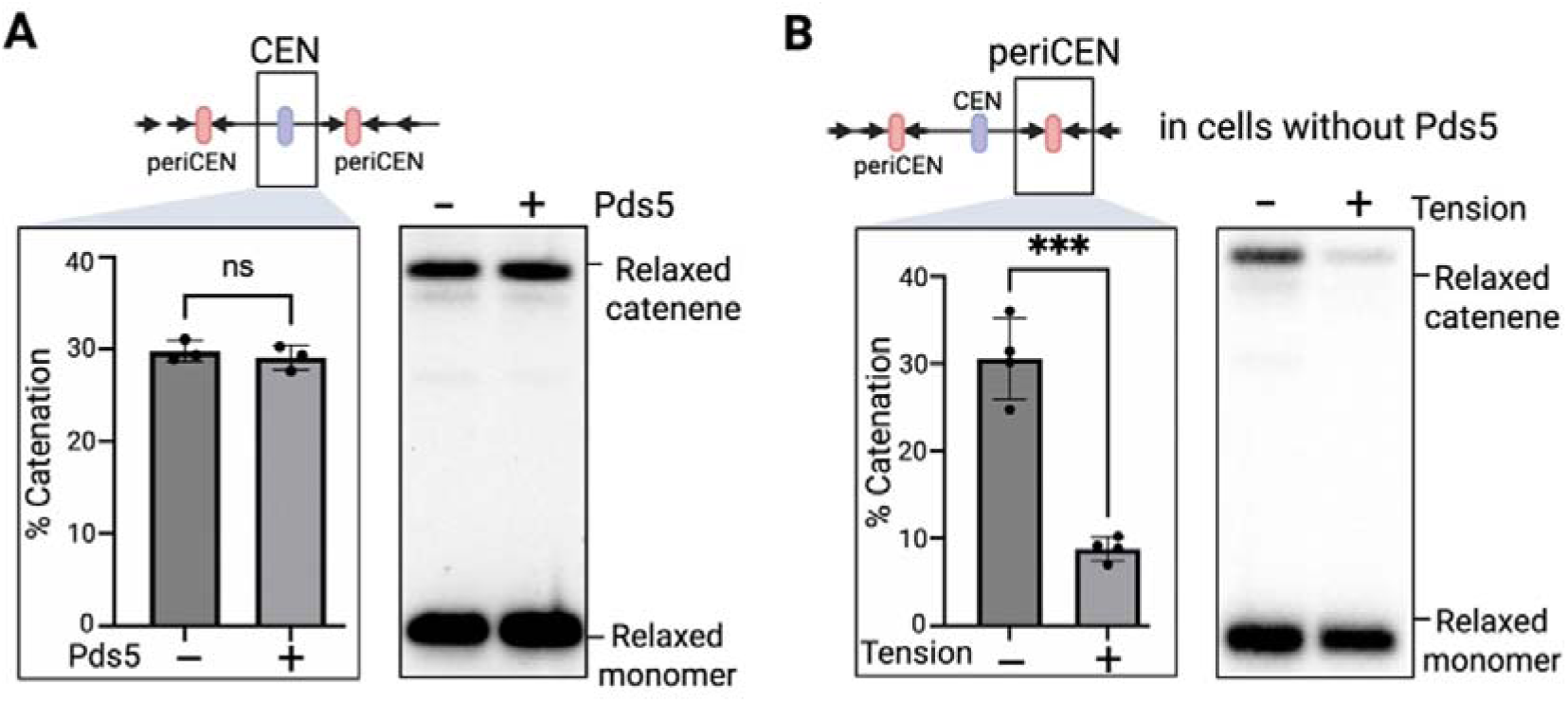
Persistence of DNA catenation does not require sister DNA co-entrapment. (A) Persistence of centromeric DNA catenation does not require sister DNA co-entrapment. Centromeric DNA catenation was measured in wild-type (29532) and *PDS5-AID* (30329) cells. Cells were synchronised in G1, Pds5 depletion was induced by auxin addition, and cells were released into mitotic arrest in the presence of nocodazole and auxin (Fig. S3B; Methods). CEN III was excised by Cre recombinase, genomic DNA was isolated, nicked with Nb.BsmI and analysed by Southern blotting. Representative Southern blots and quantification of DNA catenation from three independent biological experiments are shown. Data are mean ± s.d. **(B) Sister DNA co-entrapment is required to retain pericentromeric DNA catenation under spindle tension.** Pericentromeric DNA catenation was measured in *PDS5-AID* cells (31060) treated as in (A) and released into metaphase in the presence or absence of spindle tension (Fig. S3C; Methods). Peri-centromeric DNA was excised by Cre recombinase, genomic DNA was isolated, nicked with Nb.BsmI and analysed by Southern blotting. Representative Southern blots and quantification of DNA catenation from three independent biological experiments are shown. Data are mean ± s.d.

### Sister DNA co-entrapment anchors catenations at peri-centromeres under spindle forces

Having established that sister DNA co-entrapment is dispensable for preserving DNA catenations in cells lacking spindle forces, we next examined whether this was also true under spindle tension. In metaphase-arrested wild-type cells, we had observed that DNA catenations accumulate within pericentromeric regions under spindle tension (Fig. 1C). Unexpectedly, although depletion of Pds5 had little effect on pericentromeric DNA catenation in nocodazole-arrested cells, it abolished this tension-dependent accumulation of DNA catenations following Cdc20 depletion (Fig. 5B, Fig. S3C). These observations indicate that sister DNA co-entrapment is not required to protect DNA catenations from Topo II-mediated resolution but is essential to prevent their displacement from pericentromeric regions under spindle-generated pulling forces.

If failure to retain DNA catenations under spindle tension underlies the severe cohesion defect caused by Pds5 depletion, then loss of Pds5 should preferentially compromise cohesion when spindle forces are present. Consistent with this prediction, Pds5 depletion caused catastrophic loss of cohesion in metaphase-arrested cells, with nearly 95% of cells displaying complete sister chromatid separation (Fig. S3E,F). In contrast, Pds5 depletion produced only a modest cohesion defect in nocodazole-arrested cells lacking spindle forces, despite abolishing sister DNA co-entrapment (Fig. S3E,G). Because DNA catenations remain preserved in the absence of Pds5, we reasoned that they account for this residual cohesion. Consistent with this idea, selective removal of DNA catenations by PBCV-Topo II markedly increased sister chromatid separation in Pds5-depleted nocodazole-arrested cells (Fig. S3G), whereas PBCV-Topo II had little effect in wild-type cells. Thus, simultaneous loss of sister DNA co-entrapment and DNA catenation phenocopied cohesin inactivation in cells lacking spindle forces (Fig. 2A).

Together, these findings reveal two distinct cohesin activities that cooperate to generate force-resistant sister chromatid cohesion. One activity preserves DNA catenations independently of sister DNA co-entrapment, whereas the other anchors DNA catenations within pericentromeric regions under spindle tension through sister DNA co-entrapment.

### Pericentromeric DNA catenation requires Scc2

Our finding that DNA catenations are preserved independently of sister DNA co-entrapment suggested that their preservation depends on a distinct chromosome-organizing activity of cohesin. We therefore asked whether this process requires Scc2, the conserved activator of cohesin ATPase activity^24^ and an essential driver of cohesin-mediated loop extrusion^26^. We acutely depleted Scc2 from metaphase-arrested cells. Micro-C analysis revealed that Scc2 depletion caused a profound loss of centromeric and pericentromeric DNA loops (Fig. 6A and Fig. S4A), demonstrating that continuous Scc2 activity is required to maintain centromeric loop architecture during metaphase. If centromeric loop architecture underlies preservation of DNA catenation, its disruption should lead to catenation loss. Consistent with this prediction, Scc2 depletion caused a marked reduction in the DNA catenations that normally accumulate within pericentromeric regions under spindle tension (Fig. 6B).

**Fig. 6.**
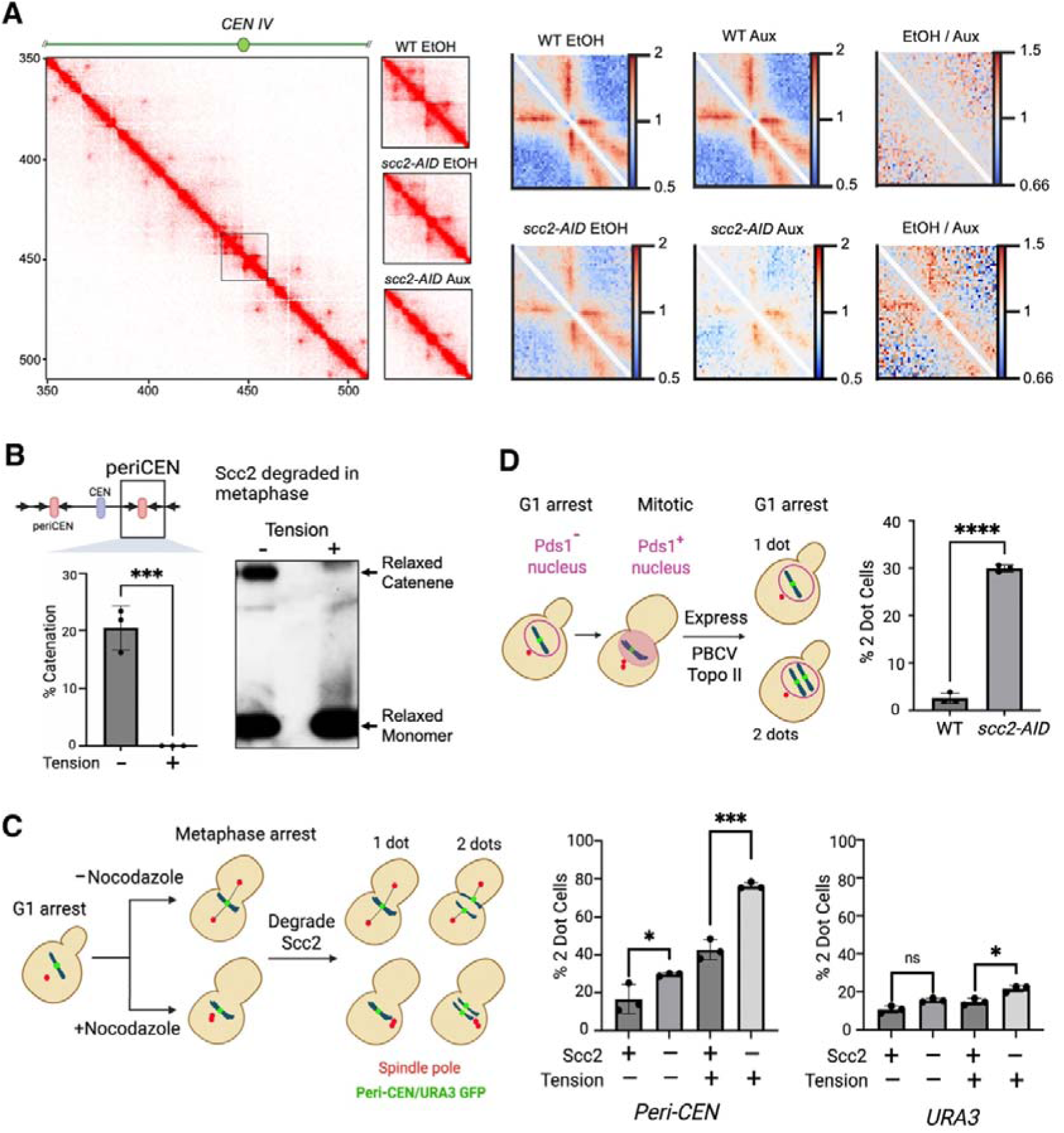
Scc2 maintains pericentromeric DNA catenation and force-resistant cohesion. (A) Scc2 maintains centromeric chromosome architecture. Micro-C contact maps (1 kb bins) of chromosome IV from wild-type (30355) *and SCC2-AID* (31684) cells treated with ethanol (EtOH) or auxin following metaphase arrest (Fig. S5A; Methods). An enlarged view of chromatin contacts surrounding *CEN IV* (black box) is shown. Pile-ups (1 kb bins) of cis contacts spanning 25 kb around all 16 centromeres and the corresponding difference maps (log₂ ratio, EtOH/Aux) highlight Scc2-dependent chromatin contacts. **(B) Scc2 maintains pericentromeric DNA catenation.** Pericentromeric DNA catenation was measured in *SCC2-AID* cells (31899) following acute Scc2 depletion in metaphase (Fig. S4B; Methods). Cells were arrested in metaphase with nocodazole, Scc2 degradation was induced by auxin addition, and peri-CEN DNA was excised by Cre recombinase. Genomic DNA was isolated, nicked with Nb.BsmI and analysed by Southern blotting. Representative Southern blot and quantification of DNA catenation from three independent biological experiments are shown. Data are mean ± s.d. **(C) Scc2 maintains force-resistant pericentromeric cohesion.** Cohesion was monitored at GFP-marked pericentromeric (Peri-CEN)(31684) and chromosome arm (URA3) loci in *SCC2-AID* cells (32632) following acute Scc2 depletion in metaphase (Fig. S4C; Methods). Cells were synchronised in G1, released into metaphase in the presence or absence of nocodazole, and Scc2 degradation induced by auxin addition. Cohesion was scored cytologically as one GFP focus (cohesed sister chromatids) or two GFP foci (separated sister chromatids). At least 100 metaphase cells were analysed per experiment. Graphs show mean ± s.d. from three independent biological experiments. **(D) Scc2 maintains faithful chromosome segregation.** Wild-type (32057) *and SCC2-AID* (32631) cells carrying GFP-marked *CEN IV* and PDS1-Myc18 were synchronised in G1 and released into metaphase in the presence of nocodazole. Scc2 was acutely depleted by auxin addition before cells were released into mitosis. Chromosome segregation was scored in G1 cells, identified by loss of nuclear Pds1 staining, as one or two GFP foci. At least 100 G1 cells were analysed per experiment. Graphs show mean ± s.d. from three independent biological experiments.

Because loss of pericentromeric DNA catenation would be predicted to impair force-resistant cohesion, we next examined sister chromatid cohesion following Scc2 depletion. Previous studies reported that Scc2 is dispensable for cohesion maintenance after metaphase establishment, based on separation of sister URA3 loci. Consistent with these observations, acute depletion of Scc2 did not detectably affect cohesion at the URA3 locus (Fig. 6C). In contrast, cohesion within the pericentromeric region was compromised (Fig. 6C). Crucially, this defect was markedly exacerbated under spindle tension(Fig. 6C). These observations indicate that Scc2 is specifically required to maintain the catenation-dependent, force-resistant cohesion associated with centromeric regions.

We next asked whether this selective loss of pericentromeric cohesion compromises chromosome segregation. Scc2 was acutely depleted in metaphase-arrested cells in the absence of spindle forces before cells were released into mitosis, and chromosome segregation was assessed in the subsequent G1 phase. Scc2 depletion resulted in a marked increase in chromosome mis-segregation, manifested by unequal inheritance of GFP-marked chromosome IV (Fig. 6D). Thus, Scc2-dependent preservation of pericentromeric DNA catenation is required for faithful chromosome segregation.

### Cohesin protects DNA knots in G1 cells from Topo II mediated resolution

Having established that cohesin preserves DNA catenations through a chromosome-organizing activity, we next asked whether this activity also protects intramolecular DNA entanglements. To address this, we examined DNA knots, intramolecular DNA entanglements that are structurally analogous to catenations between sister DNAs (Fig. S5A).

DNA knotting was quantified by high-resolution two-dimensional gel electrophoresis23 using circularised mini-chromosomes encompassing the centromeric and pericentromeric regions of chromosome II (Fig. S5B). Wild-type and *scc1-73* cells were synchronised in early G1 using α-factor at 25°C and then released into a late G1 arrest while shifting to 37°C to inactivate cohesin. Acute cohesin inactivation caused a marked reduction in DNA knotting (Fig. 7A), demonstrating that cohesin preserves both intramolecular (DNA knots) and intermolecular (DNA catenations) entanglements from Topo II-mediated resolution.

**Fig. 7.**
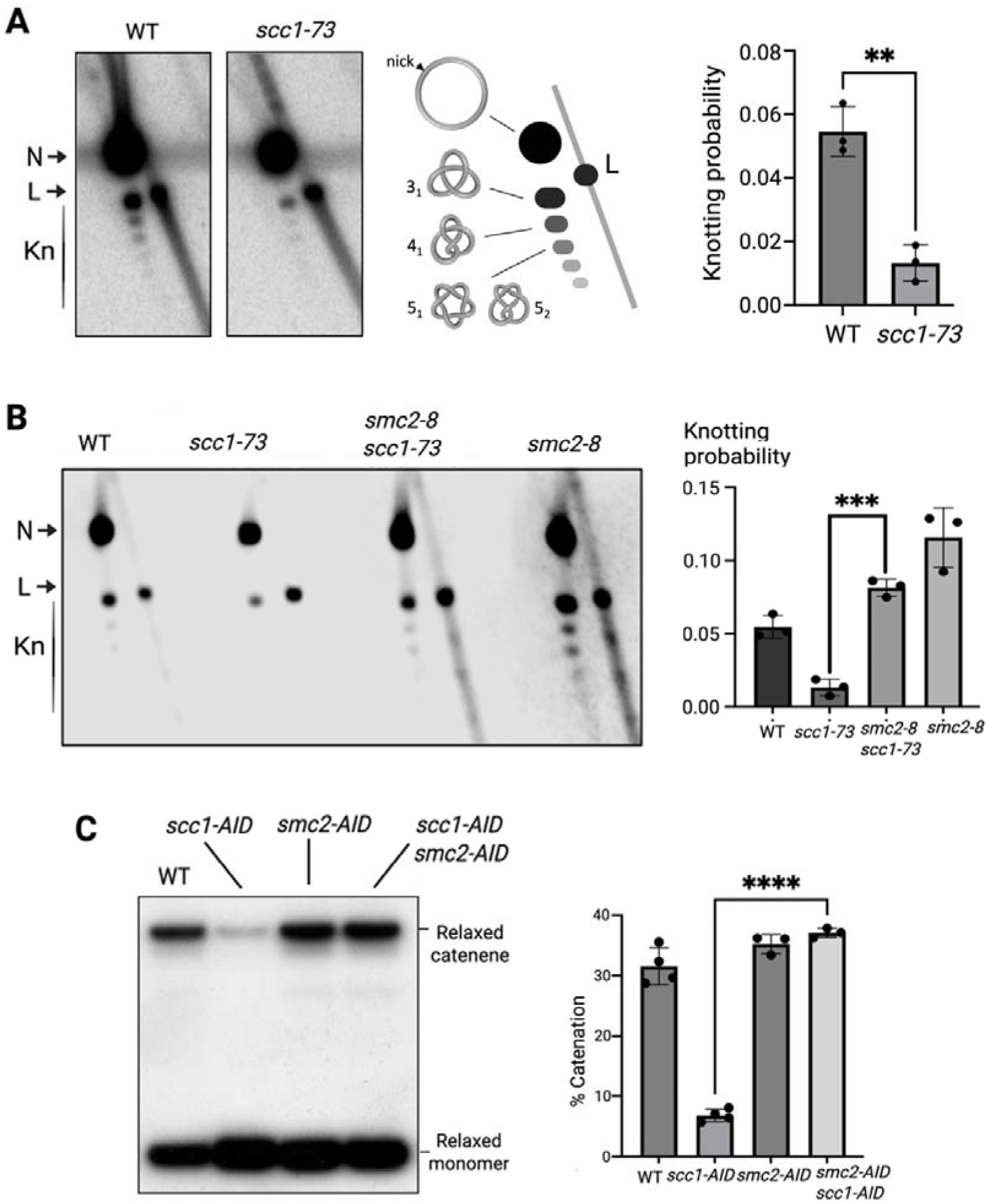
Cohesin and condensin exert opposing regulation of Topo II-mediated DNA entanglement resolution. (A) Cohesin preserves DNA knots. DNA knotting was quantified in wild-type (31457) and *scc1-73* (31553) cells carrying a circular CEN–peri-CEN II mini-chromosome. Cells were synchronised in G1 at 25°C and released into a late G1 arrest at 37°C to inactivate cohesin (Fig. S4B,C; Methods). Genomic DNA was analysed by high-resolution two-dimensional gel electrophoresis. The probability of DNA knotting was calculated from the proportion of knotted DNA relative to nicked DNA from three independent biological experiments. N, nicked; L, linear; Kn, knots of increasing complexity. **(B) Condensin promotes Topo II-mediated resolution of DNA knots.** DNA knotting was quantified in wild-type (31457), *scc1-73* (31553), *smc2-8* (31497) and *scc1-73 smc2-8* (32060) cells carrying the circular CEN–peri-CEN II mini-chromosome. Cells were synchronised in G1 at 25°C and released into a late G1 arrest at 37°C to inactivate cohesin and/or condensin (Fig. S4C; Methods). Representative two-dimensional gels and quantification of DNA knotting from three independent biological experiments are shown. **(C) Condensin promotes Topo II-mediated resolution of DNA catenations.** Centromeric DNA catenation was measured in wild-type (29532), *SCC1-AID* (30231), *SMC2-AID* (32071) and *SCC1-AID SMC2-AID* (32083) cells. Cells were synchronised in G1, depleted of the indicated AID-tagged proteins by auxin addition, and released into mitosis in the presence of nocodazole and auxin (Fig. S4E; Methods). CEN III was excised by Cre recombinase, genomic DNA was isolated, nicked with Nb.BsmI and analysed by Southern blotting. Representative Southern blots and quantification of DNA catenation from three independent biological experiments are shown. Data are mean ± s.d.

To determine whether loss of DNA knots resulted from Topo II-mediated resolution, we compared *scc1-73* single mutants with *scc1-73 top2-4* double mutants. Cells were arrested in late G1 and shifted to 37°C to inactivate cohesin and Topo II simultaneously (Fig. S5C). Topo II inactivation completely prevented the loss of DNA knots following cohesin inactivation (Fig. S5D), demonstrating that cohesin protects DNA knots by preventing their Topo II-mediated resolution.

Together with our findings on sister DNA catenations, these observations establish that protection of DNA entanglements from Topo II-mediated resolution is a general property of cohesin that extends beyond sister chromatid cohesion.

### Cohesin and condensin exert opposing effects on Topo II activity

Previous studies established that condensin is required for complete resolution of sister chromatid entanglements during anaphase ^25^, suggesting that it promotes Topo II-mediated decatenation. Consistent with this, condensin inactivation increases the steady-state level of DNA knots^27^, in striking contrast to the loss of DNA knots that we observed following cohesin inactivation (Fig. 7A). These contrasting effects suggested that cohesin and condensin exert opposing regulation of Topo II activity.

To test this idea, we first examined DNA knotting. A strain carrying the temperature-sensitive *scc1-73* and *smc2-8* alleles was used to simultaneously inactivate cohesin and condensin. DNA knotting was quantified in late G1-arrested cells by high-resolution two-dimensional gel electrophoresis. Consistent with previous observations, cohesin inactivation reduced DNA knotting, whereas condensin inactivation increased it (Fig. 7B, Fig. S5C). Strikingly, simultaneous inactivation of condensin largely suppressed the loss of DNA knots caused by cohesin inactivation (Fig. 7B), indicating that condensin is required for Topo II-mediated resolution of DNA knots even in the absence of cohesin.

We next asked whether this antagonistic regulation also extends to sister DNA catenations. To test this, we constructed a dual-degron strain enabling auxin-inducible degradation of cohesin (*Scc1-AID*) and condensin (*Smc2-AID*). Wild-type, single-degron and double-degron strains were synchronised in G1, treated with auxin and released into mitosis in the continued presence of auxin. Centromeric DNA catenation was quantified using the Cre-loxP excision assay (Fig. S5E). As observed previously, cohesin depletion markedly reduced centromeric DNA catenation, whereas condensin depletion modestly increased it (Fig. 7C). Notably, co-depletion of condensin completely suppressed the loss of DNA catenations caused by cohesin depletion (Fig. 7C), demonstrating that condensin promotes Topo II-mediated resolution of DNA catenations independently of cohesin.

Together, these findings establish opposing regulation of Topo II by cohesin and condensin as a fundamental principle of chromosome organisation. Cohesin preserves DNA entanglements by protecting them from Topo II-mediated resolution, whereas condensin promotes their removal by Topo II, revealing a general mechanism by which SMC complexes control chromosome topology.

## Discussion

Faithful chromosome segregation during mitosis depends on sister chromatid cohesion resisting microtubule-generated pulling forces long enough to generate tension and allow stable biorientation. How cells achieve this mechanical resistance has been a central question in chromosome biology for decades.

The prevailing framework for cohesion is the cohesin ring model^15^, which proposes that cohesin complexes form closed rings that topologically entrap sister DNAs following replication and maintain their association until anaphase (Fig 8A), when cleavage of cohesin by separase triggers sister chromatid disjunction^3^. This model emerged from the recognition that DNA catenation alone could not account for cohesion ^5^. Since its proposal, the ring model has explained many aspects of cohesion and has been supported by extensive experimental evidence. However, it rests on a key assumption that has never been directly tested: that topological entrapment by cohesin alone is sufficient to resist mitotic spindle forces.

**Fig. 8.**
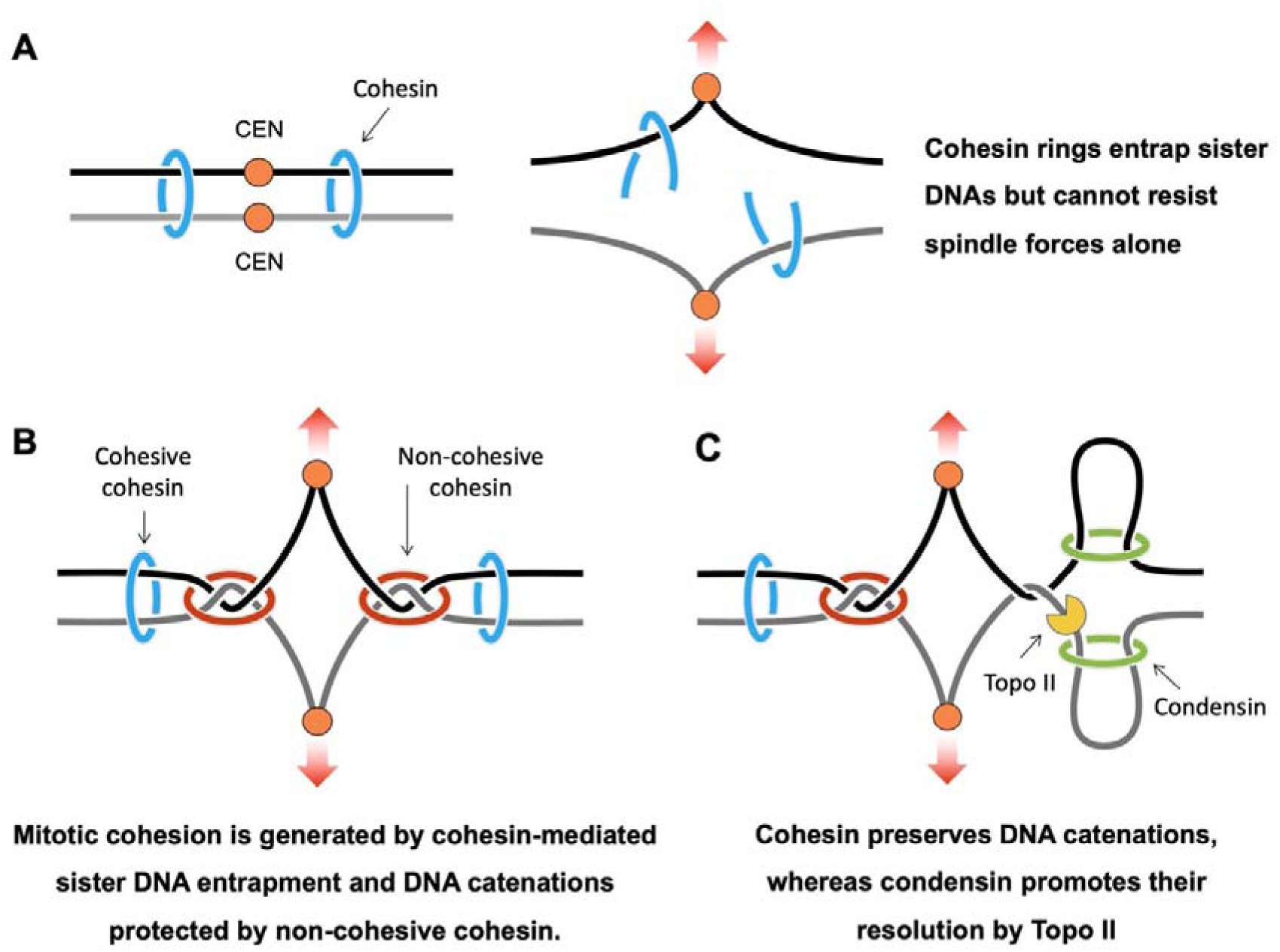
A composite topological mechanism for force-resistant sister chromatid cohesion. (A) Cohesin-mediated sister DNA entrapment alone cannot withstand spindle-generated pulling forces. Although cohesin rings topologically co-entrap sister DNAs following replication, this linkage alone is insufficient to maintain sister chromatid cohesion under spindle tension. **(B) Force-resistant sister chromatid cohesion emerges from two complementary topological mechanisms.** DNA catenations are preserved by a chromosome-organizing activity of cohesin and anchored within pericentromeric regions by cohesin rings that co-entrap sister DNAs. Together, DNA catenation and cohesin-mediated sister DNA entrapment generate a force-resistant linkage capable of resisting spindle-generated pulling forces. **(C) Cohesin and condensin exert opposing regulation of Topo II-mediated DNA entanglement resolution.** Cohesin preserves DNA entanglements by protecting them from Topo II-mediated resolution, whereas condensin promotes their removal, providing a mechanism for cell cycle-dependent regulation of DNA entanglements and timely sister chromatid separation during anaphase. Model summarizing the findings of this study. Together, these findings redefine the physical basis of sister chromatid cohesion by establishing DNA catenation as an integral, evolutionarily conserved component of the force-resistant linkage between sister chromatids.

Here we show that this assumption is incorrect (Fig 8A). Cohesin-mediated sister DNA entrapment alone cannot withstand spindle-generated pulling forces, and robust mitotic cohesion instead requires DNA catenations. Selective removal of catenations causes catastrophic loss of cohesion, delays chromosome biorientation, and increases chromosome mis-segregation, even when cohesin rings remain intact and capable of sister DNA entrapment. Conversely, persistent catenation alone cannot sustain cohesion under tension. Thus, neither cohesin-mediated entrapment nor DNA catenation alone is sufficient; mitotic cohesion emerges from their coordinated action (Fig 8B). Importantly, DNA catenation similarly underpins force-resistant cohesion in metazoans. Together with an accompanying study in human cells (Cui. R et al), our findings establish DNA catenation as an evolutionarily conserved component of the force-resistant linkage between sister chromatids.

For DNA catenations to contribute to cohesion, they must be protected from premature resolution by Topo II. Although cohesin has long been known to preserve sister chromatid catenations, this was generally attributed to cohesin-mediated sister DNA entrapment maintaining close proximity and thereby favouring re-catenation. Our findings revise this view. Preservation of catenations does not depend on co-entrapment of sister DNAs, but instead reflects a distinct activity of cohesin. The observation that cohesin similarly preserves intramolecular DNA entanglements in G1 cells indicates that protection of DNA crosses is a general property of cohesin. The requirement for Scc2 to preserve DNA catenations further suggests that this activity depends on cohesin’s chromosome-organizing functions rather than sister DNA co-entrapment (Fig 8B).

DNA catenations are also vulnerable to terminalisation by spindle forces. When tension is applied at centromeres, catenations are pushed along chromatin toward chromosome ends, progressively losing their ability to resist spindle force at centromeric regions. Our findings resolve this problem by showing that cohesin rings which co-entrap sister DNAs anchor DNA catenations within pericentromeric regions, preventing their displacement under spindle-generated pulling forces. Thus, cohesin performs two separable functions during mitosis: one chromosome-organizing activity protects DNA catenations from Topo II, whereas sister DNA co-entrapment anchors them where they are required to resist spindle force. Together, these complementary activities generate a force-resistant linkage that neither could establish alone. (Fig 8B).

Our findings also reveal how DNA entanglements are actively regulated during the cell cycle. Previous studies established that condensin promotes resolution of sister chromatid entanglements during mitosis. We show that cohesin and condensin independently exert opposing regulation of Topo II activity. Cohesin preserves DNA entanglements by protecting them from Topo II-mediated resolution, whereas condensin promotes their removal. This antagonistic regulation extends to both intramolecular DNA knots and intermolecular DNA catenations, establishing opposing regulation of Topo II by the two major SMC complexes as a fundamental principle of chromosome organisation (Fig 8C).

This antagonistic regulation provides a simple mechanism for cell cycle-dependent control of DNA entanglements. Switching cohesin off while activating condensin favours decatenation, whereas retaining cohesin preserves DNA entanglements. This logic is evident during prophase in vertebrate cells, when Wapl-mediated removal of cohesin^28,29^. coincides with condensin activation^30^ and loss of cohesion along chromosome arms, thereby reducing the risk of anaphase bridges^31^. Protection from this pathway at centromeres by shugoshins allows cohesin to preserve peri-centromeric catenations until anaphase^18^. These observations suggest that antagonistic regulation of Topo II by cohesin and condensin provides a general mechanism for controlling DNA entanglements throughout the cell cycle and raises the intriguing possibility that regulated DNA entanglements may have preceded cohesin-mediated sister DNA co-entrapment as an ancestral mechanism of chromosome cohesion.

In conclusion, our findings fundamentally redefine the physical basis of sister chromatid cohesion. Rather than being mediated solely by cohesin-mediated sister DNA entrapment, cohesion emerges from two complementary topological mechanisms. Neither element alone is sufficient: DNA catenations are vulnerable to Topo II-mediated resolution and displacement by spindle forces, whereas cohesin rings alone cannot withstand spindle tension. Together, they generate a robust force-resistant linkage that secures sister chromatids until cohesin is cleaved by separase. DNA catenation is therefore not merely a vestige of DNA replication but an evolutionarily conserved component of the force-resistant linkage that ensures accurate chromosome segregation and protects cells from aneuploidy.

## Acknowledgments

We are grateful to Kim Nasmyth and Frédéric Beckouët for stimulating discussions, and to Francis Barr, Kim Nasmyth, David Sherratt, Lars Jansen, and Frank Burmann for comments on the manuscript. We also thank the Micron Imaging Facility at the University of Oxford for technical support. Figures Created in BioRender. Srinivasan, M. (2025) https://BioRender.com/052uskt

## Author contributions

Experiments and data analysis: A.K., A.A.F., A.M.M., A.H.M., M.S. Supervision: J.R., M.S. Writing – original draft: A.K., M.S. Writing – review & editing: A.K., A.A.F., A.M.M., A.H.M., J.R., M.S. Funding acquisition: M.S.

## Data and materials availability

All data supporting the conclusions of this paper are available in the main text and/or the supplementary materials. Raw data have been deposited in Figshare *(29)*. Materials generated in this study, including plasmids and *S. cerevisiae* strains, are available upon request.

## Supplementary Materials

### Materials and methods

#### Yeast strains and plasmids

All strains are derivatives of W303 (K699). Strain numbers, plasmid numbers and relevant genotypes are listed in Supplementary Tables 1 and 2. Relevant probes and their sequences are provided in Supplementary Table 3. Antibodies and their suppliers used in the study are provided in Supplementary Table 4.

### DNA catenation assay

The yeast strain bearing the LoxP-Cre recombinase system was grown overnight in synthetic Yeast Extract Peptone (YEP), supplemented with 2% raffinose at 25 °C. The next morning, cells were diluted to 0.16-0.18 OD and synchronised in the G1 phase of the cell cycle by adding α-factor to a final concentration of 2 mg/L every 30 minutes for 2.5 hours. Cells were sedimented at 3500 rpm for 2 minutes (Multifuge 3 S-R Heraeus), and pellets were washed with 50 ml of no-sugar YEP media with 0.1 mg/ml pronase (from *streptomyces griseus*, Roche Diagnostics GmbH). Cells were released in YEP with 2% raffinose for 1.5 hours at 25 °C with 0.1 mg/ml pronase and 10µg/ml nocodazole (Sigma Aldrich) to induce mitotic arrest. Synchronisation was verified under a microscope (Zeiss AxioSkop 40) and through a Fluorescence-Activated Cell Sorting (FACS) system. Upon mitotic arrest of the strain bearing the LoxP-Cre recombinase system, 2% galactose was added and incubated at 25 °C for 1.5 hr to induce the Cre expression. Around 15OD cells were collected by centrifugation, and 25 mL of cold 50% ethanol was added to fix the cells by keeping them on ice for 30 min. For strains carrying the minichromosome, 20OD cells were collected upon mitotic arrest. Cells were sedimented, and the supernatant was discarded. The pellet was transferred to a 2 ml tube and kept frozen at 70°C till genomic DNA isolation.

### DNA knot assay

The yeast strain bearing the cen-pericen/minichromosome was grown overnight in in either YEP or Tryptophane drop out media (-Trp) supplemented with 2% raffinose at 25 °C. Next morning, cells were diluted to 0.16-0.18 OD and arrested in G1 phase of the cell cycle by adding α-factor to a final concentration of 2 mg/L, every 30 min for 2.5 hr. Cells were sedimented at 3500 rpm for 2 minutes (Multifuge 3 S-R Heraeus), and pellets were washed with 50 ml of no-sugar YEP media with 0.1 mg/ml pronase (from *streptomyces griseus*, Roche Diagnostics GmbH). Cells were released in YEP with 2% raffinose for 1.5 hours at 25 °C with 2% Galactose to induce Sic1-9m expression to synchronise cells in late G1. Synchronisation was verified under a microscope (Zeiss AxioSkop 40) and through a Fluorescence-Activated Cell Sorting (FACS) system. To evaluate the effect of temperature sensitive mutations, upon Sic1-9m mediated late G1-arrest, cells were shifted to 37°C for 90 min. Around 15OD cells were collected by centrifugation, and 25 mL of 0.5M EDTA was added to fix the cells by freezing them at-70°C.

### Yeast Genomic DNA Preparation

Yeast spheroplasting was carried out by resuspending cell pellets in 200 µl SCE buffer (1 M sorbitol, 0.1 M sodium citrate, pH 7.0, 50 mM EDTA, 0.1 M β-mercaptoethanol, 1 mg/ml zymolyase 100T from *Anthrobacter luteus*) with gentle tapping and incubating at 37°C for 1 h without shaking. Cell lysis was initiated by adding 200 µl lysis buffer (0.5 M EDTA, 0.1 M Tris-HCl pH 8.8, 10% SDS) and gently inverting tubes, followed by incubation at 65°C for ≤20 min. DNA was precipitated by adding 200 µl 5 M potassium acetate and incubating on ice for 30 min. Cell debris was pelleted by centrifugation at 13,000 rpm for 10 min at 4°C, and the supernatant was transferred to 2 ml tubes containing 200 µl 3 M sodium acetate. DNA was precipitated with 1 ml of isopropanol by gentle inversion and centrifuged at 2,300 × g for 1 min. The supernatant was carefully removed, and the pellet was washed with 700 µl of 70% Ethanol by centrifugation at 13,000 rpm for 5 min at 4°C. Supernatant was removed carefully and DNA pellets were air-dried at 37°C. Pellets were resuspended in 100 µl elution buffer supplemented with 5 µl RNase and incubated at 37°C for 2 h, without shaking.

### DNA digestion

For AfeI digestion, a reaction mix containing 2.5 µl AfeI (NEB), 2 µl 10× CutSmart buffer, and nuclease-free water to a final volume of 20 µl was prepared and added to 30 µg of DNA. Reactions were incubated at 37°C for 16 h. For linearising or nicking the looped out DNA, 30 µg of DNA was incubated with 2.5 µl SmaI (NEB) at 37°C for 4 h, or with 2.5 µl nicking enzyme Nb.BsmI (NEB) at 65°C for 4 h. For DNA knot assay, 50ugs of DNA were nicked with Nt.BspQI (NEB) for 4h at 50°C with shaking (1000 rpm), stopped by adding DNA loading buffer and heating at 65°C for 5min, then concentrated by SpeedVac.

### Differential Sedimentation of minichromosome

After synchronising cells in G2 using Nocodazole as explained before, 20 OD cells were harvested at 3500 rpm using a Heraeus Multifuge and washed twice with cold MQ H₂O. Pellets were resuspended in 100 mM Tris-HCl (pH 9.4) containing 10 mM DTT and 10 μg/ml nocodazole, then incubated on ice for 20 min. After washing with ice-cold H₂O, cells were resuspended in spheroplasting buffer (1 M sorbitol, 0.1 M sodium citrate, pH 7.0, 50 mM EDTA, 0.1 M β-mercaptoethanol, 1 mg/ml zymolyase 100T) and incubated for 30 min at 4°C with shaking. Spheroplasts were pelleted using a Beckman Coulter JA25.50 rotor at 6000 rpm for 6 min, gently washed with 1 M sorbitol, transferred to 1.5 ml tubes, and sedimented again at 1500 × g for 1 min at 4°C. Pellets were resuspended in 200 μl of cold 0.4 M sorbitol and lysed on ice for 30 min by adding 700 μl of lysis buffer (25 mM HEPES-KOH [pH 8], 50 mM KCl, 10 mM MgSO₄, 0.25% Triton X-100, 1 mM PMSF, 3 mM DTT, and 1× EDTA-free protease inhibitor cocktail), supplemented with 100 μg/ml RNase A and 300 mM NaCl. Cell extracts were obtained by centrifuging at 12,000 × g for 5 min at 4°C. Cleared lysates (450 μl) were layered onto sucrose gradients prepared using a Biocomp gradient station and centrifuged in an SW41 rotor (Beckman Optima L-100 XP preparative ultracentrifuge) at 18,000 rpm for 4 h. Gradients were fractionated using a Gilson FC203B fractionator, collecting 15 drops per fraction.

### Southern blot analysis

For DNA catenation, digested DNA was separated on a 0.8% Agarose gel made with TAE buffer containing 1.4 µg/ml Ethidium Bromide at 1V/cm for 22hr. For minichromosome sedimentation assay, gradient fractions were separated on 0.8% agarose gels containing 5 μg/ml ethidium bromide at 1.1 V/cm for 44 or 60 hr at 4°C. For the DNA knots assay, TBE buffer was used to prepare two-dimensional gel slabs, and a 2% agarose base layer (16×16cm) was first cast on the gel tray. A 0.5 cm thick resolving layer containing 2% agarose was then poured on top of this base. For two-dimensional analysis, the wells in the resolving slab were kept sufficiently apart. Electrophoresis was performed in 1 L gel tanks with an effective electrode distance of 22 cm. In the first dimension, DNA samples were separated at 25V for 42 hours at room temperature. After rotating the gel by 90°, the second dimension was conducted in the same TBE buffer at 110V for 4 hours at room temperature. DNA was transferred to a Nylon membrane (Amersham™ Hybond™-XL) following the standard alkaline transfer method. Blots were hybridised with a centromeric/pericentromeric/Trp1 probe (see Probes section for specific sequence) and scanned in FLA-7000IR (Fuji Film) and analysed using Aida Image Analyzer v4.22.

### Quantification of Southern blot

All experiments were repeated 3 times. Band intensities were measured using Aida Image Analyzer v4.22 and the following formulae were used to calculate percentages of catenation/loop out/DNA knots:

*% of catenation=Intensity of relaxed catenene band/(Intensities of relaxed catenene+ relaxed monomer bands)*

*% Of loopout= Intensity of post looped out DNA band/(Intensities of pre+ post looped out DNA bands)*

*% Of DNA knots= Intensity of DNA knot band/(Intensities of DNA knots+ nicked plamid bands)*

### *In vivo* BMOE crosslinking

Yeast strains containing a 2.3 Kb minichromosome were grown in Tryptophane dropout medium at 25°C to an OD600 of 0.5–0.6. Next morning, cells were synchronised in G1 phase of the cell cycle by adding α-factor to a final concentration of 2 mg/L, every 30 min for 2.5 hr. Synchronised cultures in G1 were washed with YEP-no sugar and released into YPAD containing 8 mM methionine; methionine was replenished every hour at 4 mM to maintain repression. After 1.5 hr, 30 OD units of cells were harvested, washed with ice-cold PBS, and resuspended in 500 µl ice-cold PBS and 30 µl BMOE (42 mg/ml stock in DMSO). Samples were incubated on ice for 6 min, then washed twice with 2 ml ice-cold PBS and kept frozen at-70°C till lysis.

### Minichromosome immunoprecipitation

Pellets were resuspended in 500 µl lysis buffer (25 mM HEPES pH 8.0, 50 mM KCl, 50 mM MgSO₄, 10 mM trisodium citrate, 25 mM sodium sulfite, 0.25% Triton X-100) freshly supplemented with Complete Protease Inhibitor (Roche, 2×) and PMSF (1 mM). Cells were lysed using a FastPrep-24 (MP Biomedicals) for 3 × 1 min at 6.5 m/s with 500 µl acid-washed glass beads (425–600 µm, Sigma), and lysates were clarified by centrifugation at 12,000 × g for 5 min. Protein concentrations were adjusted after Bradford assay and cohesin immuno-precipitated using anti-PK antibody (AbD Serotec, 2 h, 4°C) and protein G dynabeads (2 h, 4°C, with rotation). After cohesin immunoprecipitation, protein G Dynabeads were washed twice with 1 ml lysis buffer, resuspended in 30 µl 1% SDS containing DNA loading dye, and incubated at 65°C for 4 min. Supernatants were resolved on 0.8% agarose gels containing ethidium bromide (1.4 V/cm, 22 h, 4°C). Following alkaline Southern blot transfer, DNA was detected using a [³²P]-labeled TRP1 probe.

### Sister chromatid cohesion assay

Cells containing pMET-CDC20 were synchronised in G1 in methionine-free synthetic medium following the same procedure as described earlier. Cells were then arrested in metaphase in the presence or absence of spindle tension as described in Paldi et al., 2020. Briefly, synchronised cultures in G1 were washed with YEP-no sugar and released into YEPR containing 8 mM methionine; methionine was replenished every hour at 4 mM to maintain repression. Samples were collected after 90 minutes, and for each repeat, at least 50 cells were scored for GFP dots. For metaphase arrest in the absence of microtubule-dependent tension, cells were released from G1 as mentioned earlier but with Nocodazole (10 µg/ml). Samples were collected after 90 minutes, and for each repeat, at least 50 cells were scored for GFP dots. For temperature-sensitive mutants, after mitotic arrest, cells were shifted to 37°C and incubated for 90 mins.Samples were collected after 90 minutes, and for each repeat, at least 50 cells were scored for GFP dots.

For chromosome segregation assay, to arrest cells lacking the pMET-CDC20 construct in metaphase without spindle tension, asynchronous cultures (OD600 = 0.2) were treated with nocodazole (15 µg/ml) and benomyl (30 µg/ml), with nocodazole replenished every hour at 7.5 µg/ml. Cells were harvested after 3 h. Cells were washed with rich media (YPAD) and released into pre-warmed YPDA at 30°C containing α-factor to a final concentration of 2 mg/L. Cells were harvested after 90 min and fixed using formaldehyde as previously described. Cells with negative Pds1-Myc18 were scored for one or two GFP dots.

For biorientation assays in fixed cells, cells were first arrested without tension in YEPR media containing nocodazole (15 µg/ml) and benomyl (30 µg/ml) for 1.5 hr and 2% Galactose was added upon Mitotic arrest to express PBCV-Gal TopII. After another 1.5 hr, nocodazole and galactose was removed by centrifugation, and cells were washed with rich medium containing 2% glucose (YPAD) before releasing into YPDA+Met to allow spindle reformation while maintaining metaphase arrest. Samples were collected every 20 min, and for each repeat, at least 50 cells were scored for GFP dots.

For biorientation assays in live cells, after removing nocodazole and galactose by centrifugation, and cells were washed with SC-complete medium containing 2% glucose. Cells were then placed on SC-complete Agarose medium containing Methionine, and imaging was done in 3-minute intervals for 1.5 hr. Around 60 cells were tracked for GFP dots separation following SPB separation.

### Microscopy of fixed cells

To count the GFP dots, cells were fixed with 4% formaldehyde for 1 hour at room temperature, washed 2 times with 1x PBS and stored at 4°C in PBS-Sorbitol (1xPBS+1M sorbitol). Cells were placed on a glass slide, and GFP dots were counted by fluorescence microscopy using Zeiss Axio Imager Z1 microscope equipped with a 63×/1.40 NA objective and a CoolSNAP HQ camera. All dot counting experiments were repeated 3 times, and at least 50 cells were analysed for each repeat. The GFP dots were scored by double blinding.

### Live cell microscopy

Cells were placed on SC-complete agarose medium containing 4 mM methionine and images were recorded in at least four well-separated locations, every 3 min for 90 min. Images were acquired with a Leica THUNDER system (Leica) equipped with a DMi8 inverted motorised stand, a 63x HC PL APO CS2 1.20 NA water automated immersion objective and an ORCA-fusion digital CMOS camera (Hamamatsu), using 470 nm (GFP) and 542 nm (tdTomato) lasers from a Coolled pE-800 LED Light Engine (127 W). Time lapses were taken of cells in an incubation chamber set at 30 °C. 3D image stacks were obtained in z with 20-30 raw images per position per time point. Adaptive focus control (AFC) corrected for z-stack drift over the time-lapses. For image analysis, Iif files were opened on ImageJ to produce z-stack maximum projections that were corrected for photobleaching via histogram matching. In all experiments, there was about a 20-minute delay between the placement of the cells on the agarose pad and the start of imaging. n=60 cells were tracked for each strain.

### I*n situ* immunofluorescence detection of Pds1-MYC

Fixed cells were spheroplasted with Zymolyase 100T for 30 min at 30°C. Spheroplasts were immobilised on poly-L-lysine–coated slides and permeabilised with 1% NP-40 for 5 min. Slides were blocked in PBS containing 1% BSA (PBS-BSA) and incubated overnight at 4°C with anti-MYC (Sc-40 Anti MYC from Santacruz) antibody diluted in PBS-BSA. After ten washes in PBS-BSA, slides were incubated with fluorescently labelled AlexaFluor 647 mouse secondary antibody (Thermo Fisher Scientific) for 2 h at room temperature. Samples were mounted in DAPI-containing mounting medium and imaged using a Zeiss Axio Imager Z1 microscope equipped with a 63×/1.40 NA objective and a CoolSNAP HQ camera. All experiments were performed in triplicate, and at least 50 Pds1 negative cells were analysed per replicate.

### Auxin inducible degradation of Scc1/Pds5/Smc2 proteins

Cultures were arrested in G1 for 1.5 hr, as explained in previous sections before adding 10mM Indole-3-acetic acid sodium salt (Chemcruz) and incubating for another 1 hr. Cells were released to mitotic/metaphase arrest in the presence of 10mM Indole-3-acetic acid sodium salt and incubated for another 1.5 hours before processing for the next steps, depending on the experiment.

### Auxin inducible degradation of Scc2

Cells containing pMET-CDC20 were synchronised in G1 in methionine-free synthetic medium followed by arrested in metaphase in the presence or absence of spindle tension as described earlier. Scc2 was degraded using 10mM Indole-3-acetic acid sodium salt (Chemcruz) and incubating for another 1 hr. Cells were collected, fixed for microscopy and GFP dots were counted as explained before.

For chromosome segregation assay, cells lacking pMET-CDC20 were arrested in absence of tension and Scc2 was degraded as mentioned earlier. Cells were then washed with rich media (YPAD) and released into pre-warmed YPDA at 30°C containing α-factor to a final concentration of 2 mg/L. Cells were harvested after 90 min and fixed using formaldehyde as previously described. Cells with negative Pds1-Myc18 were scored for one or two GFP dots.

### Western blot detection of proteins

20OD cells were sedimented by centrifugation to isolate the protein and carry out a western blot to examine degradation of the targeted protein/s. Cell pellet/s was lysed in 700 µl lysis buffer (25 mM HEPES pH 8.0, 50 mM KCl, 50 mM MgSO₄, 5 mM β-Mercaptoethanol, 0.5% NP-40 0.25% Triton X-100) freshly supplemented with Complete Protease Inhibitor (Roche, 2×) and PMSF (1 mM). Lysis was performed using a FastPrep-24 instrument (MP Biomedicals) for 3 × 1 min at 6.5 m/s with 500 µl acid-washed glass beads (425–600 µm, Sigma). Lysates were clarified by centrifugation at 13,000 × g for 10 min. Protein concentrations were determined using the Bradford assay. Whole-cell lysates were separated on NuPAGE 4–12% gradient gels (NuPAGE® Life Technologies) and transferred to PVDF membranes using the Trans-Blot Turbo transfer system (Bio-Rad). Membranes were probed with appropriate antibodies (Table S4), followed by incubation with Immobilon Western chemiluminescent HRP substrate (Millipore). Signals were detected using either an ODYSSEY Fc Imaging System (LI-COR) or X-ray exposure.

### Microinjection

Microinjection experiments were performed as previously described in Oliveira et al., 2010; Piskadlo et al., 2017. Briefly, 1-1.5 hr old dechorionated embryos were glued to a #1.5 coverslip, covered with Series 700 halocarbon oil (H8898; Sigma-Aldrich) and injected at 18–20°C into the posterior pole using a Burleigh Thorlabs micromanipulator, a Femtojet microinjection system (Eppendorf), and prepulled Femtotip I needles (Eppendorf). Injections were performed using 9.8 mg/ml PBCV1 Top2 protein diluted in 20 mM Tris-HCl, pH 7.5, 300 mM NaCl, 6 mg/ml TEV protease in TEV buffer (20 mM Tris-HCl, pH 8.0, 1 mM EDTA, 50 mM NaCl, and 2 mM DTT).

### Live cell imaging of Drosophila embryo

LIce cell imaging was performed on an inverted widefield DeltaVision microscope (Applied Precision Ltd.) at 18–20°C in a temperature-controlled room using a 100× oil-immersion 1.4 NA objective lens (Olympus), and standard live filter sets. Time lapses were taken every minute with z series optical sections recorded every 0.8 µm with SoftWoRx software (5.5.0; Applied Precision Ltd.). Widefield images were restored by conservative deconvolution with SoftWoRx software and were assembled using FIJI software.

### Top2 purification

Top 2 purification was carried out as described in Srinivasan et al. 2019. Briefly, Top2 bearing a Strep-II tag was purified from 500 ml of SF-9 insect cells, infected with baculovirus stock in a 1/100 dilution. Cells were harvested when lethality reached no more than 70–80%. Cell pellets were then frozen in liquid nitrogen and stored at 80°C. Upon thawing, the pellets were suspended in a final volume of [65–70 ml with Buffer A (25 mM HEPES pH 8.0, NaCl 150 mM, TCEP-HCl 1 mM and Glycerol 10%), supplemented with two tablets of Roche Complete Protease (EDTA-free), 75 μg of RNase I and 7 μl of DNaseI (Roche, of 10 U/μl stock). To lyse the cells, sonication was done at 80% amplitude for 5 s/burst/35 ml of suspension using a Sonics Vibra-Cell (3 mm microtip). In total, 12 bursts were given for every 35 ml. 235,000 x g, 45 min centrifugation was done (45,000 rpm on a Ti45 fixed angle rotor) following addition of PMSF to 1 mM final concentration. The cleared extract was supplemented with 2 mM EDTA and loaded to a 2 × 5 ml StrepTrap HP (Fisher Scientific) column at 1 ml/min in an ÄKTA Purifier 100. The column was washed with Buffer A at 1 ml/min to the point of ΔΑU280nm[0 and protein elution was carried out using Buffer A + 20 mM desthiobiotin (Fisher Scientific) at 1 ml/min. Peak fractions were analysed using SDS-PAGE and further purified in a Superose 6 Increase 10/300 (VWR) using Buffer A as running buffer (free of EDTA/PMSF). The resulting peaks were quantified in Nanodrop and analysed using SDS-PAGE.

### Micro-C library preparation

The Micro-C protocol was adapted from Nguyen et al^32^ with minor modifications. Cells were fixed with 3% formaldehyde for 15 min at 30°C. Cross-linking was quenched by adding glycine to a final concentration of 0.2 M, followed by incubation with shaking for 5 min at 30°C. Cells were harvested, washed with sterile ice-cold water, and subjected to cell-wall permeabilization by resuspension in 10 mL of buffer Z (1 M sorbitol, 50 mM Tris-HCl pH 7.5, 10 mM 2-mercaptoethanol) containing 100 μg/mL Zymolyase 100T (Nacalai Tesque, 07665-55). Samples were incubated for 15 min at 30°C with inversion every 5 min. Spheroplasts were washed with ice-cold PBS, transferred to Costar 1.7 mL Low Binding Snap Cap Microcentrifuge Tubes (Corning, 3207), pelleted by centrifugation, and resuspended in 1 mL PBS containing 3 mM DSG (Thermo Fisher Scientific, 20593) for a second cross-linking step. Samples were incubated for 40 min at 30°C with shaking at 125 rpm, with tube inversion every 10 min. Cross-linking was quenched by adding glycine to a final concentration of 0.4 M and incubating for 5 min at 30°C with shaking. Cells were washed with ice-cold PBS, pelleted by centrifugation, snap-frozen in liquid nitrogen, and stored at-80°C.

For chromatin fragmentation, frozen cell pellets were resuspended in 500 μL of ice-cold MB1 (50 mM NaCl, 10 mM Tris-HCl pH 7.5, 5 mM MgCl₂, 1 mM CaCl₂, 0.005% NP-40, 0.5 mM spermidine, 1.43 mM 2-mercaptoethanol, 1× cOmplete EDTA-free protease inhibitor cocktail (Roche, 04693132001)) and incubated on ice for 5 min. After centrifugation, pellets were resuspended in 500 μL MB1 containing 250 U MNase (NEB, M0247S) and incubated for 20 min at 37°C with shaking at 1250 rpm. Reactions were terminated by adding EGTA to a final concentration of 2.5 mM, followed by incubation at 65°C for 10 min. Fragmented chromatin was pelleted and washed twice with ice-cold MB2 (50 mM NaCl, 10 mM Tris-HCl pH 7.5, 10 mM MgCl₂) before end preparation.

A three-step end-preparation reaction was performed as previously described^33,34^. The reaction was terminated by adding EDTA to a final concentration of 30 mM followed by incubation at 65°C for 20 min. Chromatin was washed once with 1× T4 DNA ligase buffer (NEB, B0202S) and pelleted by centrifugation. For proximity ligation, chromatin pellets were resuspended in 500 μL ligation mixture containing 1× T4 DNA ligase buffer, 0.1 mg/ml recombinant albumin (NEB, B9200S), and 5,000 CEU T4 DNA ligase (NEB, M0202L), and incubated on a slow rotator for 3 h at room temperature. Biotin-dNTPs remaining at unligated DNA ends were removed by incubation with 400 U Exonuclease III (NEB, M0206S) in 1× NEB buffer 1 (NEB, B7001S) for 15 min at 37°C. Chromatin was then incubated with 200 μg/mL RNase A (VWR, A3832) in TE containing 0.5% SDS for 1 h at 50°C, followed by incubation with 3 mg/mL Proteinase K (Sigma, 03115879001) and 1% SDS for 2 h at 55°C and overnight incubation at 65°C. DNA was purified using a QIAquick PCR Purification Kit (QIAGEN, 28106) and eluted in buffer EB (QIAGEN, 19086). An initial end repair was performed using 50 μL of purified DNA with the NEBNext Ultra II DNA Library Prep Kit for Illumina according to the manufacturer’s instructions.

Biotin-labeled DNA fragments were captured by incubation with 5 μL Dynabeads MyOne Streptavidin C1 beads (Invitrogen, 65001) in binding buffer (BB; 5 mM Tris-HCl pH 7.5, 0.5 mM EDTA, 1 M NaCl) on a slow rotator for 20 min at room temperature. Beads were washed twice with 200 μL Tween washing buffer (TWB; 5 mM Tris-HCl pH 7.5, 0.5 mM EDTA, 1 M NaCl, 0.05% Tween-20), washed once with 200 μL buffer EB, and resuspended in 50 μL buffer EB. Library preparation was then carried out on the beads using the NEBNext Ultra II DNA Library Prep Kit for Illumina from the end-repair step through adapter ligation according to the manufacturer’s instructions.

Following adapter ligation, beads were washed twice with 200 μL TWB and once with 200 μL buffer EB followed by resuspension in 15 μL buffer EB. PCR amplification was performed using the NEBNext Ultra II DNA Library Prep Kit for Illumina according to the manufacturer’s instructions. PCR products were purified using SPRIselect (Beckman Coulter, B23318), followed by size selection for fragments between 260 and 575 bp using SPRIselect. Libraries were eluted in 15 μL buffer EB and sequenced as paired-end reads on an Illumina NextSeq 1000.

### Micro-C library analysis

The Micro-C data were processed using CustardPy v2.2.0 with the default parameter^35^. The sequenced reads were mapped to the S. cerevisiae genome obtained from Saccharomyces Genome Database. The uniquely mapped read pairs were randomly resampled and arranged in the same numbers within sample groups. Contact matrices used for further analysis were background-SQRT normalized at 1kb resolution with Juicer^36^. The matrices were visualized by Juicebox^37^.

Pile-up plots of pixels at centromeres/pericentromeric borders were calculated and normalized to the expected signal of global cis interactions at 1-kb resolution. The centromeres/pericentromeric borders were described in Paldi et al. Plots were created around the midpoints of centromeres with 25-kb flanks on each side, or around the midpoint of borders with 25-kb flanks. All centromere/pericentromere annotations were duplicated in the forward and reverse strand orientations to create an image that is mirror symmetrical. Briefly, cool format files were obtained using the cload pairix algorithm and the obtained matrices were normalized using the balance algorithm in cooler version 0.9.3^38^. Expected signals for cis interactions were calculated from these matrices using the expected-cis algorithm in cooltools version 0.7.0^39^. Pile-up plots were then calculated using coolpup.py and plotted using plotpup.py in coolpup.py version 1.1.0^40^.

### Statistics and reproducibility

Sample size was not pre-determined. Sample size and the number of biological replicates for each experiment are indicated in the figure legends. Student’s t-test was used for determining the statistical significance. Mean, SD and statistical significance are shown in each figure. No data were excluded from the analyses.

---

**Fig S1.**
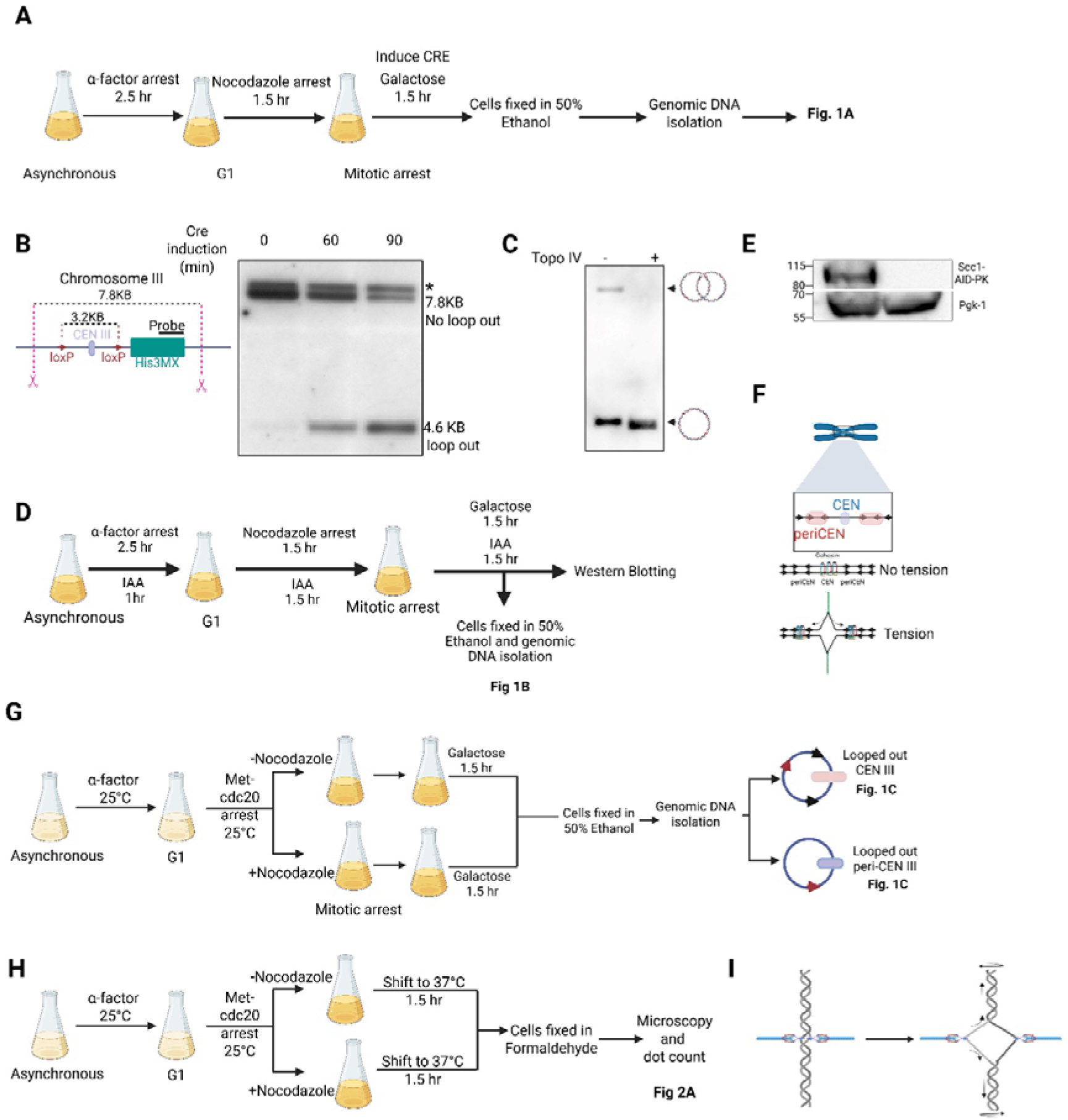
(A) Exponentially growing strain 29532 in YEP + raffinose medium was arrested in G1 using α-factor for 2.5 h, then released into a mitotic arrest induced by nocodazole. Cre recombinase expression was induced by adding galactose (2% final concentration) for 90 min. Cells were fixed with cold 50% ethanol and genomic DNA was isolated as described in *Methods*. **(B)** Efficiency of CEN III excision by CRE induction. The Cre recombinase was induced in cells arrested in mitosis, as described in (A), for the indicated times. Genomic DNA was isolated, digested with AfeI, and subjected to agarose gel electrophoresis and southern blotting with a probe complementary to the *HIS3 MX* cassette. * denotes a background band seen in all lanes arising from the endogenous *HIS3* gene. See *Methods* for details about the southern blot probes and the method of quantification. **(C)** The genomic DNA from (A) was nicked with Nb-BsmI and treated with *E.coli* Topo IV at 37°C for 60 minutes. Digested DNA was analysed by agarose gel electrophoresis and Southern blotting using CEN probes. **(D)** Experimental setup for Fig. 1B. Exponentially growing wild-type strain 29532 and *SCC1-AID* strain 30245 were arrested in G1 using α-factor. Depletion of Scc1 was initiated by the addition of IAA. The cultures were released into a mitotic arrest in media containing both auxin and Nocodazole. CRE recombinase expression was induced for 90 minutes, and DNA was isolated from fixed cells. **(E)** Western blot for the culture described in (D) examining depletion of Scc1 with reference to Pgk1 as loading control. Whole cell extracts were loaded onto 3-8% gradient gels and subjected to SDS-PAGE and western blotting (See *Methods)*. The western blots were probed with the indicated antibodies. **(F)**Diagram illustrating cohesin relocation from centromeres (CEN) to peri-centromeric regions (peri-CEN) under spindle tension *(18)*. **(G)** Experimental setup for Fig. 1C. Exponentially grown strains 30411 and 30904 were arrested in G1 using pheromone α-factor in YEP Raffinose medium. Cells were released into YEP Raffinose medium with 8 mM Methionine to repress the expression of Cdc20, in the presence and absence of Nocodazole. Galactose was added for 90 mins to induce CRE recombinase in the metaphase-arrested cells. Cells were fixed, and genomic DNA was isolated according to the protocol described in *Methods* **(H)** Experimental setup for Fig. 1D. Exponentially grown strains 31001 and 31044 were arrested in G1 using pheromone α-factor in YEP glucose medium. Upon G1 arrest, cells were released into a metaphase arrest induced by Methionine repressible *CDC20* in the presence and absence of Nocodazole at 25°C. The cultures were then shifted to restrictive temperature 37°C for 90 min, and microscopy was performed on formaldehyde-fixed cells as described in *Methods.* **(I)**Spindle forces will displace catenations along the chromosome arms towards chromosome ends.

**Fig S2.**
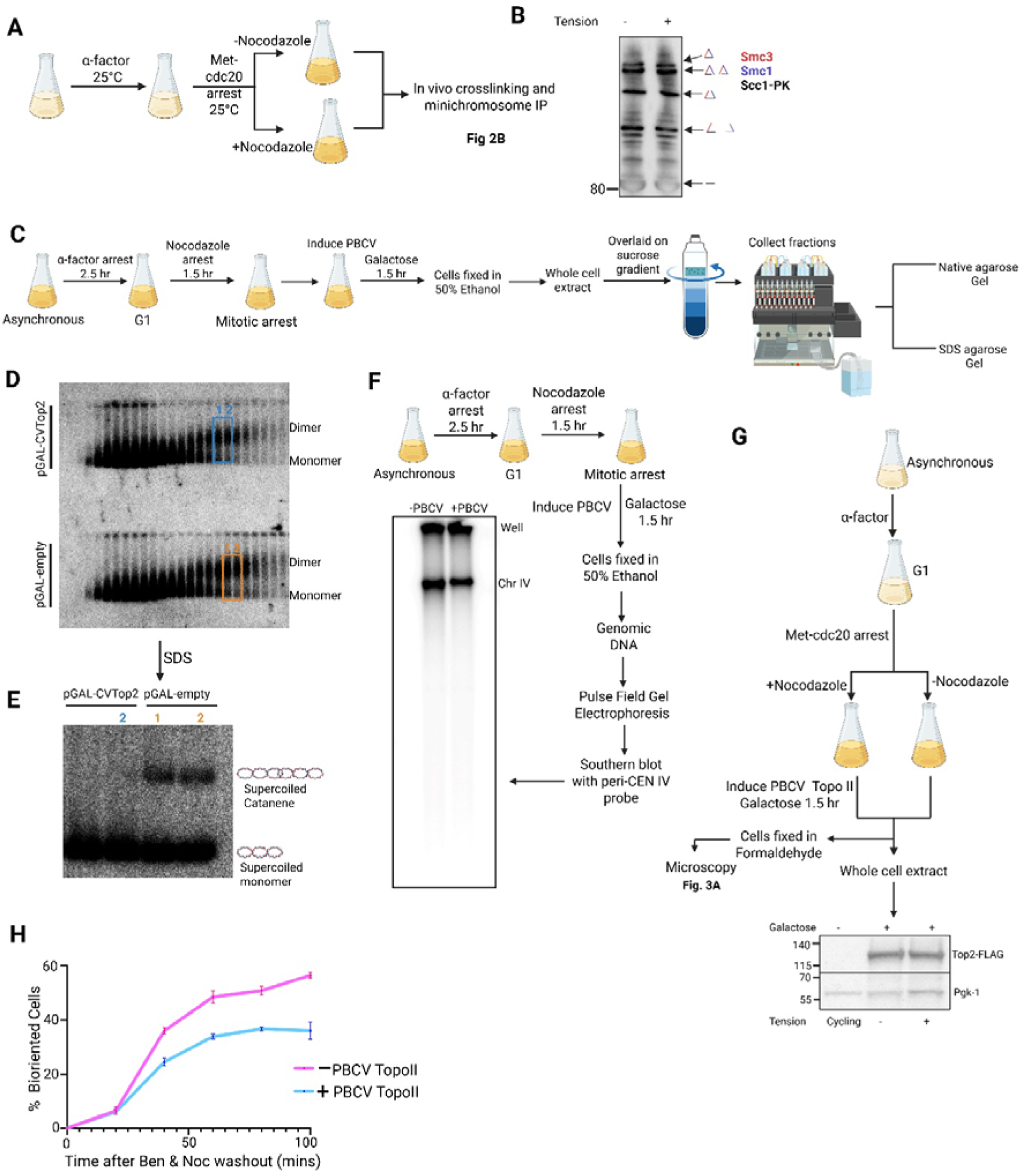
(A) Experimental set up for Fig 2A. Exponentially growing strain 32191 in methionine tryptophan dropout medium was arrested in G1 with α-factor. Synchronised cells were released into metaphase arrest induced by Methionine repressible *CDC20* in the presence and absence of Nocodazole. *In vivo* BMOE crosslinking followed by minichromosome immunoprecipitation was performed on metaphase arrested cells, as described in *Methods* **(B)** Western blot showing Scc1-immunoprecipitation for the strain 32191. Whole cell extracts were immunoprecipitated using anti-V5 antibody against Scc1-PK. Samples were loaded onto 3-8% gradient gels and subjected to SDS-PAGE and western blotting. See *Methods* for immunoprecipitation and western blotting details. **(C)** Experimental setup for (D) and (E). Exponentially growing strain 16150 in YEP with raffinose medium was arrested in G1 using α-factor and released into a mitotic arrest induced by nocodazole. PBCV-Top2 expression was induced with galactose for 90 min in mitotically arrested cells. Whole-cell extracts from fixed cells were separated on a sucrose gradient; fractions were collected and analyzed by native and SDS agarose gel electrophoresis. See *Methods* for details on gel conditions and Southern blotting. **(D)** Native agarose gel from the experiment described in (A), showing that dimers can be detected in both the presence and absence of PBCV-Top2. **(E)** SDS agarose gel for the experiment explained in (A) showing PBCV-TopoII expression resolved catenated minichromosomes. **(F)** Experimental setup for pulsed-field gel electrophoresis showing the absence of non-specific chromosome breaks upon PBCV-Top2 induction. Exponentially growing strain 30548 in YEP medium with raffinose was arrested in G1 using α-factor pheromone and released into a mitotic arrest induced by nocodazole. Overexpression of PBCV-Top2 was induced for 90 min by adding galactose to mitotically arrested cells. DNA was extracted from fixed cells and analysed by pulsed-field gel electrophoresis as described in *Methods*. **(G)**Experimental setup for Fig. 2B. Exponentially growing strain 30548 was subjected to G1 arrest induced by α-factor pheromone. Upon arrest, cells were released into a metaphase arrest induced by Methionine repressible *CDC20* in the presence or absence of Nocodazole. PBCV Top2 overexpression was induced by Galactose for 90 mins. Cells were collected, fixed, and microscopy was performed as described in *Methods*. Western blot showing overexpression of PBCV-Top2 protein with reference to Pgk1 as loading control in the whole cell extracts of cells described in (A). See M*ethods* for whole cell extract preparation and western blot conditions. **(H)**Fixed cell microscopy for cells shown in C. Cells were collected every 20 min after releasing from Nocodazole and Benomyl arrest till 100 min. For every time point, ≤50 cells were imaged as described in *Methods*. Every time point is repeated for three times. See methods for microscopy details.

**Fig S3.**
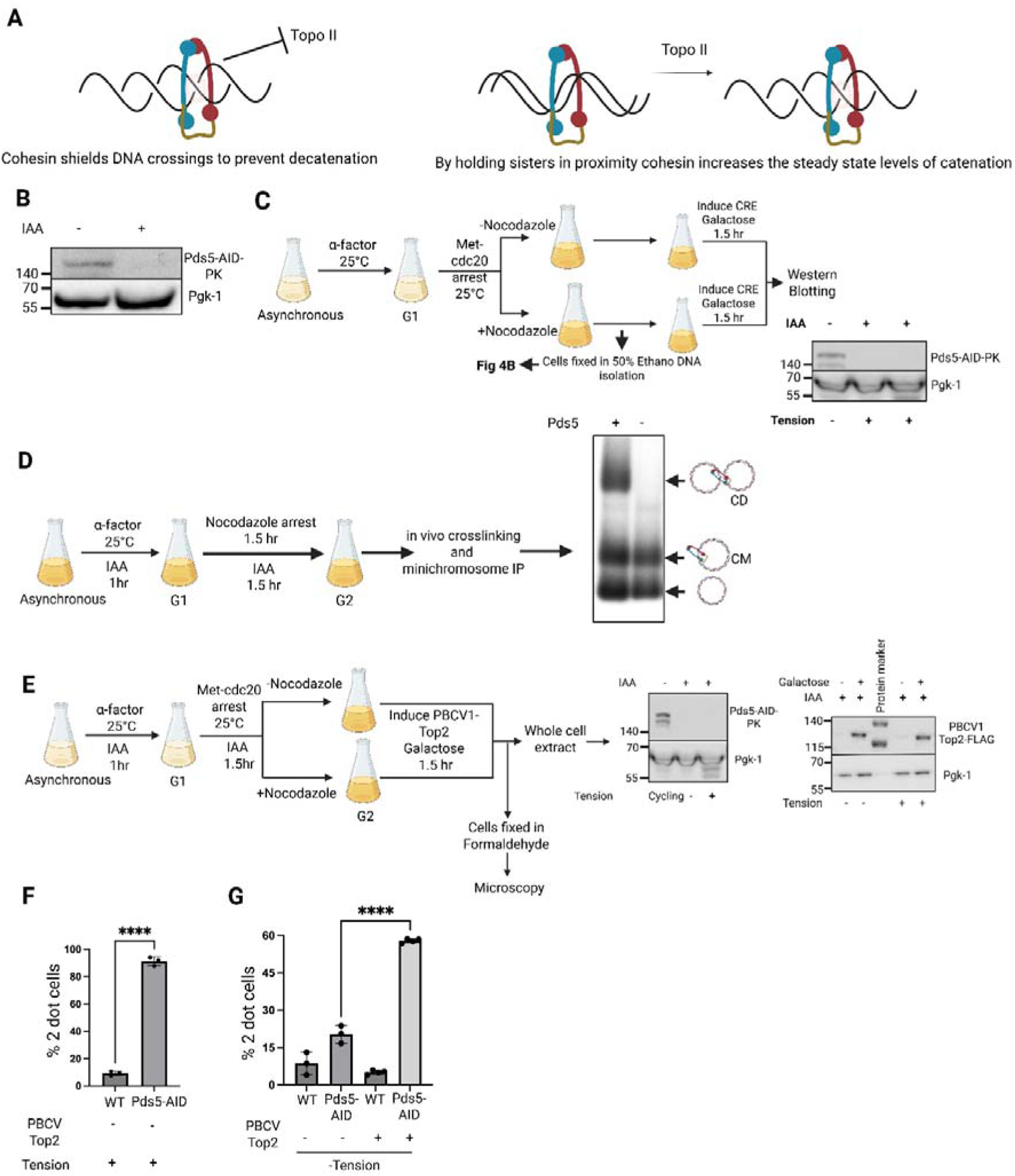
(A) Possible models for how cohesin preserves DNA catenation—either by shielding DNA crossings from Top2-mediated decatenation or by holding sister chromatids in close proximity to maintain catenation equilibrium (see Results). **(B)** Western blot for Fig. 3A. Whole-cell extracts were analysed to confirm Pds5 depletion, with Pgk1 as a loading control as explained in the methods section. **(C)** Experimental setup for Fig. 3B. Exponentially growing strain 31060 was arrested in G1 with α-factor. Pds5 depletion was initiated by adding IAA after 1.5 h in α-factor–containing medium. Following G1 arrest, cultures were released into a methionine-repressible CDC20–dependent metaphase arrest in the presence of auxin, with or without nocodazole. Samples for Western blotting were collected before Cre recombinase expression was induced in metaphase-arrested cells by adding galactose for 90 min. DNA was isolated from fixed cells, and Southern blotting was performed using a probe against pericentromere III. Whole-cell extracts were separated on 4–12% gradient gels and probed for Pds5 depletion, with Pgk1 as a loading control. See methods for details **(D)** Experimental setup and results for analysis of sister DNA co-entrapment after Pds5 depletion. Exponentially growing strain 31807 in tryptophan dropout medium was arrested in G1 with α-factor. Pds5 depletion was initiated by adding IAA after 1.5 h in α-factor–containing medium, and cultures were released into nocodazole-induced mitotic arrest in the presence or absence of auxin. In vivo BMOE crosslinking followed by mini-chromosome immunoprecipitation was performed on mitotically arrested cells, as described in Methods **(E)** Experimental setup for (F) and (G). Strain 32117 grown in YEP + raffinose medium was arrested in G1 with α-factor, and Pds5 depletion was initiated by adding IAA after 1.5 h. Following G1 arrest, cultures were released into a methionine-repressible CDC20–dependent metaphase arrest in the presence of auxin, with or without nocodazole. Samples were collected for Western blotting before galactose was added for 90 min to induce PBCV-Top2 expression. Cells were fixed and analysed by microscopy, and Western blotting confirmed Pds5 depletion, with Pgk1 as a loading control as described in *Methods*. **(F)** ≥100 metaphase cells were scored for one or two GFP dots in WT and Pds5-aid cells in the presence of tension, and each experiment was repeated three times. **(G)** ≥100 metaphase cells were scored for one or two GFP dots in WT and Pds5-aid cells in the absence of tension and presence or absence of overexpressed PBCV Top2. Each experiment was repeated three times.

**Fig. S4.**
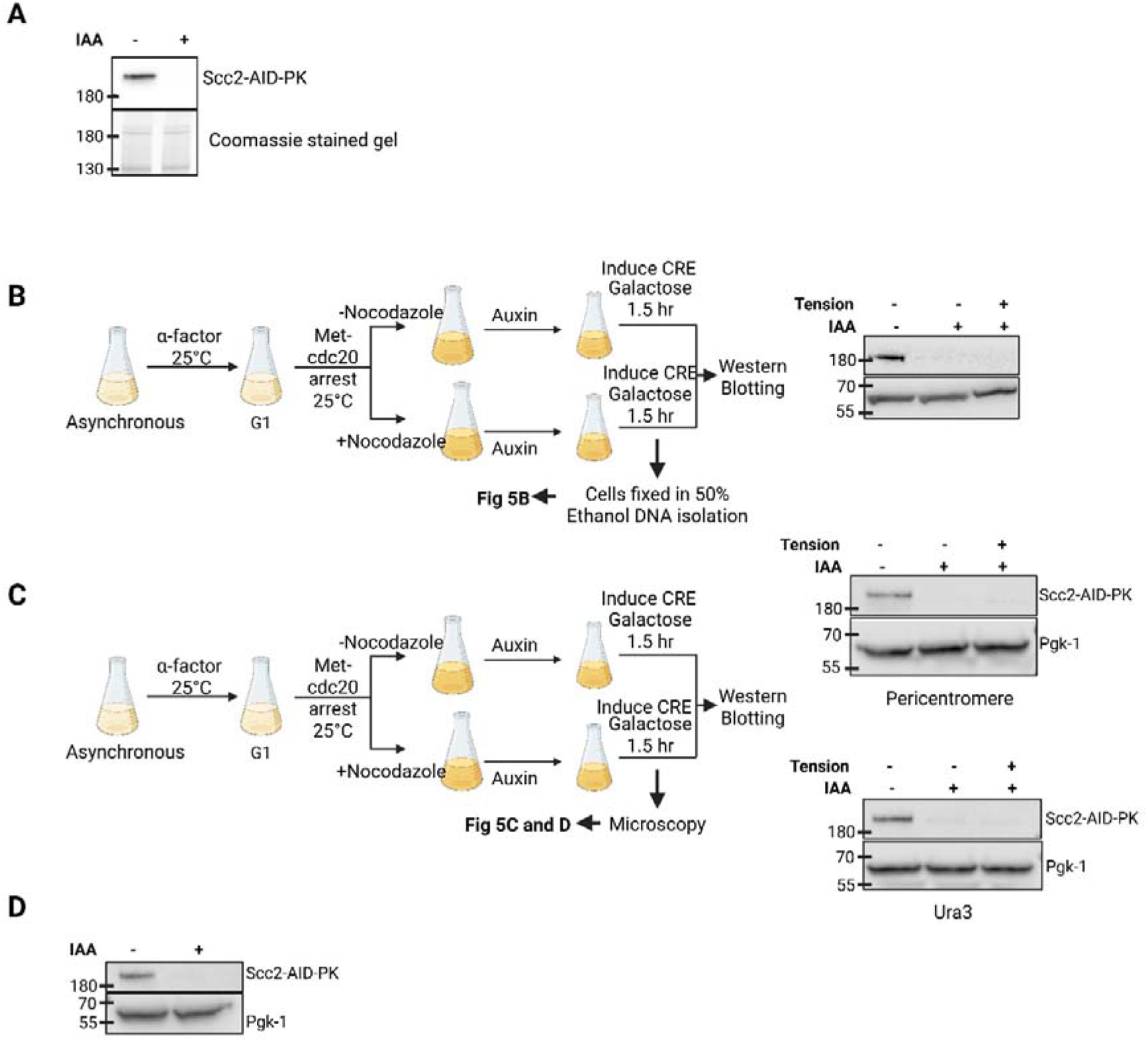
(A) Western blot confirming Scc2 depletion, with Coomassie-stained gel as control**(B)** Experimental setup for Fig. 6B. Exponentially grown strain 31899 was arrested in G1 using pheromone α-factor in YEP Raffinose medium. Cells were released into YEP Raffinose medium with 8 mM Methionine to repress the expression of Cdc20, in the presence and absence of Nocodazole. Metaphase arrested cells were treated with Auxin and Galactose was added for 90 mins to induce CRE. Cells were fixed, and genomic DNA was isolated according to the protocol described in Methods. Western blotting confirmed Scc2 depletion, with Pgk1 as a loading control. **(C)** Experimental setup for Fig. 6C. Strains 32634 and 31684 were grown in YPD medium was arrested in G1 with α-factor. Following G1 arrest, cultures were released into a methionine-repressible CDC20–dependent metaphase arrest in the presence of auxin, with or without nocodazole. Upon arrest, Scc2 depletion was initiated by adding auxin for 1.5hr. Samples were collected for Western blotting and cells were fixed and analysed by microscopy. Western blotting confirmed Scc2 depletion, with Pgk1 as a loading control as described in Methods. (D) Western blot confirming Scc2 depletion, with Pgk1 as a loading control for Fig. 6D.

**Fig S5.**
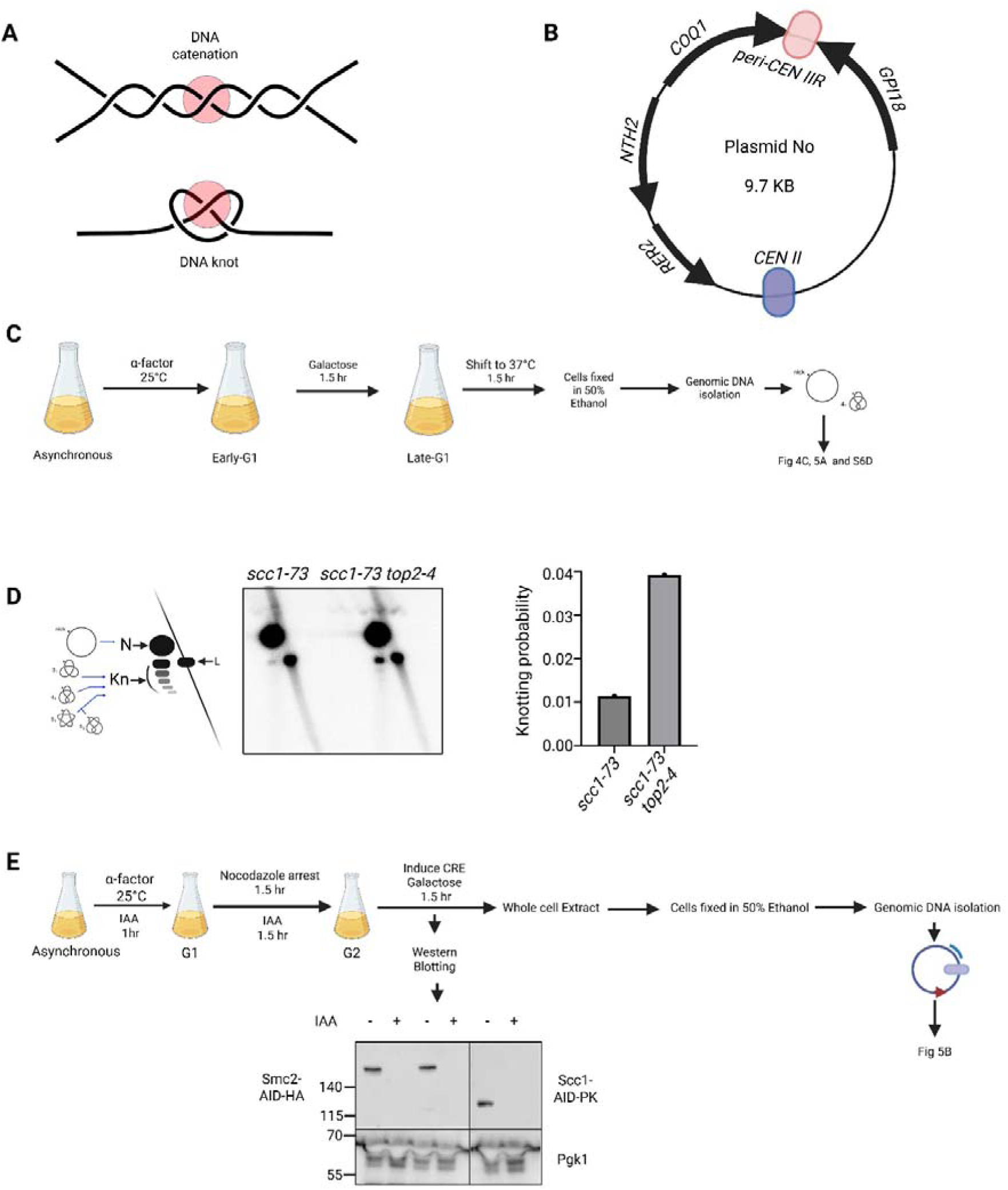
(A) Schematic illustrating that DNA crossings in catenations and knots are topologically equivalent **(B)** Schematic map of the plasmid carrying the centromeric (CEN) and pericentromeric (peri-CEN) regions of chromosome II used in Fig. 3C. **(C)** Experimental setup for Fig. 4A. Exponentially growing strains WT (31457), *scc1-73* (31553), *smc2-8* (31497), and *scc1-73 smc2-8* (32060) were arrested in G1 with α- factor pheromone and then released into a late G1 arrest by overexpressing non-degradable Sic1-9m. Cultures were shifted to the non-permissive temperature for 90 min. DNA was isolated from fixed cells, and Southern blotting was performed as described in the Methods. DNA bands were detected using a probe against pericentromere II. **(D)** The extent of DNA knotting in a circular CEN–peri-CEN II mini-chromosome was analysed in *scc1-73* (31553) and *scc1-73 top2-4* (31556) cells. Cells were synchronized in G1 at 25°C and released into a late G1 arrest at 37°C to inactivate cohesin (Fig. S6C; *Methods*). Genomic DNA was resolved by high-resolution two-dimensional gel electrophoresis, and the proportion of knotted DNA relative to nicked plasmid was quantified from three independent experiments. N, nicked; L, linear; Kn, knots of increasing complexity **(E)** Experimental setup for Fig. 4A. Exponentially growing strains WT (29532) *SCC1-AID* (30245) *SMC2-AID* (32071) and *SMC2*-AID *SCC1-*AID (32083), in YEP with raffinose medium were arrested in G1 with α-factor. Depletion of Scc1-AID and Smc2-AID was initiated by the addition of IAA after 1.5 h of α-factor treatment. Cultures were then released into mitotic arrest in medium containing both auxin and nocodazole. Mitotically arrested cells were collected for whole-cell extract preparation before Cre recombinase expression induction by adding galactose. Western blotting was performed using antibodies against Smc2-HA and Scc1-PK, with Pgk1 as a loading control to examine the depletion of Scc1 and Smc2. DNA isolation and Southern blotting were performed as described in *Methods*, using a probe against centromere III.

**Table S1:** Yeast strains used in this study.

| **Strain Number** | **Genotype** |
| --- | --- |
| **16150** | Mat a, 26kb minichromosome (TRP1, ARS1, CEN3 ~22KB) |
| **29532** | Mat a, Smc1(G22C,K639C)::NatMX4, Smc3(E570C,S1043C)::ADE2, LoxP CEN III LoxP::HIS3MX, ura3::Gal CRE::URA3, CEN III flanked by LoxP, LoxP upstream of YCL001W-A |
| **30245** | Mat a, Smc3(E570C,S1043C)::ADE2, SCC1-PK3-AID::KanMX4, his3::ADH1 promoter-OsTIR1-9myc::HIS3LoxP CEN III LoxP::HIS3MX, ura3::Gal CRE::URA3 |
| **30329** | Mat a, PDS5-Pk3-aid::KanMX4, his3::ADH1 promoter-OsTIR1-9myc::HIS3LoxP CEN III LoxP ::HIS3MX, ura3::Gal CRE::URA3, CEN III flanked by LoxP |
| **30411** | Mat a, MET3-CDC20::TRP1, his3::ADH1 promoter-OsTIR1-9myc::HIS3, LoxP CEN III LoxP ::HIS3MX, ura3::Gal CRE::URA3, CEN III flanked by LoxP |
| **30484** | MAT a, MET-CDC20::URA3, Spc42-tdTomato::NAT, leu2::PURA3::tetR-GFP::LEU2, tetOx224-URA3 (tetOs~11.5kb to right of CEN4, OUTSIDE the boundary), pRS423::His |
| **30548** | MAT a, MET-CDC20::URA3, Spc42-tdTomato::NAT leu2::PURA3::tetR-GFP::LEU2, tetOx224-URA3 (tetOs~11.5kb to right of CEN4, OUTSIDE, the boundary), Gal-PBCV-1 Top2::His |
| **30904** | MAT a, ura3::Gal CRE::URA3Met-cdc20::leu, loxP pericenIII loxP::Trp |
| **31001** | MAT a, scc1-73::HIS MX, MET-CDC20::URA3, Spc42-tdTomato::NAT, leu2::PURA3::tetR-GFP::LEU2, tetOx224-URA3 (tetOs~11.5 kb to right of CEN4, OUTSIDE the boundary) |
| **31023** | MAT a, PDS5-Pk3-aid::KanMX4, his3::ADH1 promoter-OsTIR1- |
|  | 9myc::HIS3ura3::Gal CRE::URA3, loxP pericen III loxP::Trp |
| **31044** | MAT a, scc1-73::HIS MX, Top2-4::Trp, MET-CDC20::URA3, Spc42-tdTomato::NAT, leu2::PURA3::tetR-GFP::LEU2, tetOx224-URA3 (tetOs~11.5 kb to right of CEN4, OUTSIDE the boundary) |
| **31060** | MAT a, PDS5-Pk3-aid::KanMX4, his3::ADH1 promoter-OsTIR1-9myc::HIS3, Met-cdc20::leu, ura3::Gal CRE::URA3, loxP pericen III loxP::Trp |
| **31457** | MAT a, leu2::Gal1p-Sic1(9m)/His3p-Gal1/His3p-Gal2/Gal1p-Gal4::Leu2 (single copy), Cen2 PeriCen2 TRP1 ARS1 10.9 kb Plasmid |
| **31497** | MAT a, smc2-8, leu2::Gal1p-Sic1(9m)/His3p-Gal1/His3p-Gal2/Gal1p-Gal4::Leu2 (single copy), Cen2 PeriCen2 TRP1 ARS1 10.9 kb Plasmid |
| **31553** | MAT a, Scc1(S525N)::His3MX6 (scc1-73), leu2::Gal1p-Sic1(9m)/His3p-Gal1/His3p-Gal2/Gal1p-Gal4::Leu2 (single copy), Cen2 PeriCen2 TRP1 ARS1 10.9 kb Plasmid |
| **31556** | MAT a, smc2-8, leu2::Gal1p-Sic1(9m)/His3p-Gal1/His3p-Gal2/Gal1p-Gal4::Leu2 (single copy), Cen2 PeriCen2 TRP1 ARS1 10.9 kb Plasmid |
| **31684** | MAT a, MET-CDC20::URA3, Spc42-tdTomato::NAT, leu2::PURA3::tetR-GFP::LEU2, tetOx224-URA3 (tetOs~11.5kb to right of CEN4, OUTSIDE the boundary), Scc2-AID-PK3::KanMX, his3::ADH1 promoter-Os TIR1-9Myc::HIS3 |
| **31807** | MAT a, PDS5-Pk3-aid::KanMX4, Scc1(A547C)-pk6::Kan,Smc3 (E570C,S1043C)::ADE2, Smc1(G22c,K639C)::Nat, ura::ADH1 promoter-OsTIR1-9myc::URA3, Trp1 Ars1-cen4 2.3KB plasmid |
| **31899** | MAT a, MET-CDC20:: leu2, Scc2-AID-PK3::KanMX, his3::ADH1 promoter-OsTIR1-9Myc::HIS3, ura3::Gal CRE::URA3, loxP pericen III loxP::Trp |
| **32020** | MAT a, Spc42-tdTomato::NAT, leu2::PURA3::tetR-GFP::LEU2, tetOx224-URA3 (tetOs ~4 kb to right of CEN4, INSIDE the boundary), Pds1myc18::TRP(*K.lactis*), ADE2, pRS423::His |
| **32021** | MAT a, Spc42-tdTomato::NAT, leu2::PURA3::tetR-GFP::LEU2, tetOx224-URA3 (tetOs ~4 kb to right of CEN4, INSIDE the boundary), Pds1myc18::TRP(*K.lactis*), ADE2, Gal-PBCV1Top2::His |
| **32060** | MAT a, smc2-8, Scc1(S525N)::His3MX6 (scc1-73), leu2::Gal1p-Sic1(9m)/His3p-Gal1/His3p-Gal2/Gal1p-Gal4::Leu2 (single copy), Cen2 PeriCen2 TRP1 ARS1 10.9 kb Plasmid |
| **32057** | MAT a, MET-CDC20::URA3, leu2::PURA3::tetR-GFP::LEU2, Spc42-tdTomato::NAT, Pds1myc18::TRP(K.lactis), tetOx224-URA3 (tetOs~4 kb to right of CEN4, INSIDE the boundary), ADE2 |
| **32071** | MAT a, Smc2-HA6-AID::Hyg, his3::ADH1 promoter-OsTIR1-9myc::HIS3, LoxP CEN III LoxP::HIS3MX, ura3::Gal CRE::URA3, CEN III flanked by LoxP, LoxP Upstream of YCL001W-A and downstream of YCR001W:HIS3MX |
| **32073** | MAT a, MET-CDC20::URA3, leu2::PURA3::tetR-GFP::LEU2, Spc42-tdTomato::NAT, Pds1myc18::TRP(K.lactis), tetOx224-URA3 (tetOs~4 kb to right of CEN4, INSIDE the boundary), ADE2, pRS423::His3 |
| **32074** | MAT a, MET-CDC20::URA3, leu2::PURA3::tetR-GFP::LEU2, Spc42-tdTomato::NAT, Pds1myc18::TRP(K.lactis), tetOx224-URA3 (tetOs~4 kb to right of CEN4, INSIDE the boundary), ADE2, Gal-PBCV1Top2::His |
| **32083** | MAT a, SCC1-PK3-AID::KanMX4, Smc2-HA6-AID*::Hyg, his3::ADH1 promoter-OsTIR1-9myc::HIS3, LoxP CEN III LoxP ::HIS3MX, ura3::Gal CRE::URA3, CEN III flanked by LoxP, LoxP Upstream of YCL001W-A and downstream of YCR001W :HIS3MX |
| **32117** | MAT a, PDS5-Pk3-aid::KanMX4, his3::ADH1 promoter-OsTIR1-9myc::HIS3, MET-CDC20::URA3, leu2::PURA3::tetR-GFP::LEU2, tetOx224-URA3 (tetOs~11.5 kb to right of CEN4, OUTSIDE the boundary), Spc42-tdTomato::NAT, Gal-PBCV1-Top2::Trp |
| **32191** | MAT a, Met-cdc20::Leu, Scc1(A547C)-PK6::KAN, Smc3 (E570C,S1043C)::ADE2, Smc1(G22C,K639C)::NAT, Trp1 Ars1-cen4 2.3 kb plasmid |
| **32631** | MAT a, Spc42-tdTomato::NAT, leu2::PURA3::tetR-GFP::LEU2, tetOx224-URA3 (tetOs ~4 kb to right of CEN4, INSIDE the boundary), Pds1myc18::TRP(K.lactis), ADE2, Scc2-AID-PK3::KanMX, his3::ADH1 promoter-Os TIR1-9Myc::HIS3 |
| **32632** | Mat a, MET-CDC20:: leu2, Scc2-AID-PK3::KanMX, his3::ADH1 promoter-OsTIR1-9Myc::HIS3, tetR-GFP:LEU, TetOx200:URA3 |

**Table S2:** Plasmids generated in this study.

| Plasmid | genotype |
| --- | --- |
| **3957** | MET3-CDC20::TRP1 |
| **4493** | MET3-CDC20::LEU2 |
| **7683** | Gal-PBCV1Top2::His |
| **7687** | Cen2 PeriCen2 TRP1 ARS1 10.9 kb |
| **7691** | Gal-PBCV1Top2::Trp |

**Table S3:** Probes generated in this study.

| Probe | Sequence |
| --- | --- |
| **Cen III** | YCL001W-YCR001W |
| **Pericen III** | YCR007-YCR008 |
| **Trp1** | Trp1 gene from *S. cerevisiae* |
| **CEN-Pericen II** | Not I digestion of p7687 |
| **His3** | His3 MX |

**Table S4:** Antibodies used in this study.

| Antibody | Provider |
| --- | --- |
| **Anti PK** | Anti-V5 tag, BioRad |
| **Anti HA** | Anti HA high affinity, Roche |
| **Anti FLAG** | Monoclonal ANTI-FLAG<sup>®</sup> M2, Sigma Aldrich |
| **Anti-m9ouse for PK/FLAG** | Anti-Mouse IgG, F(ab)<sub>2</sub>, Abcam |
| **Anti-RAT for HA** | Goat anti-Rat IgG, Merck |
| **Sc-40 Anti MYC** | Santacruz |
| **AlexaFluor 647 mouse secondary antibody** | Thermo Fisher Scientific |

**Table S5:**
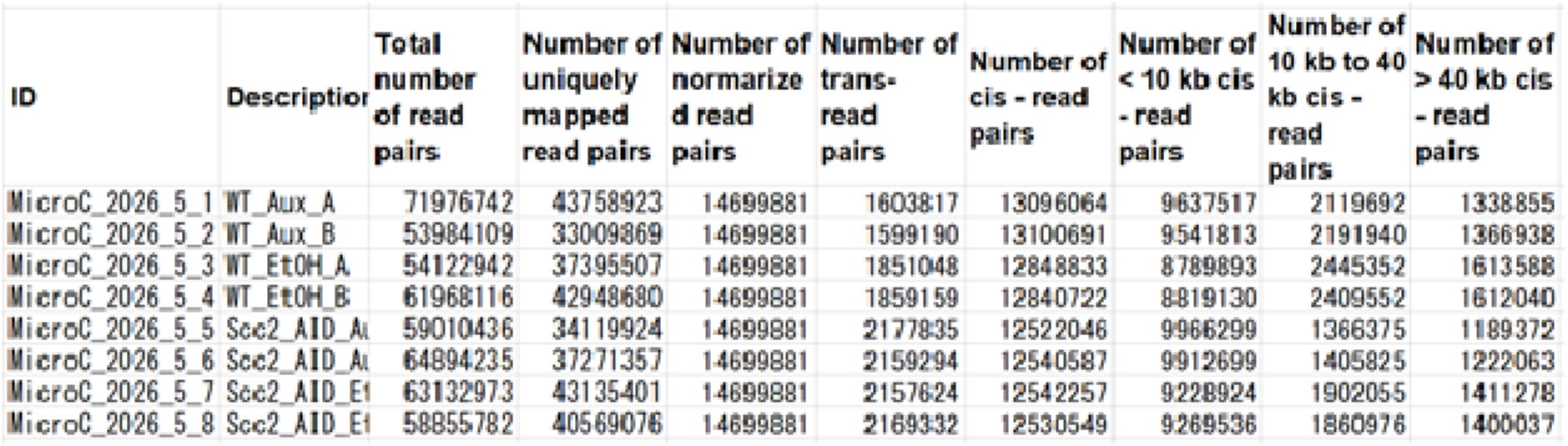
Micro-C sequencing statistics.

